# Enabling automatic generation of protein-ligand complex datasets with atomistic detail

**DOI:** 10.64898/2026.01.15.699426

**Authors:** Torben Gutermuth, Emanuel S. R. Ehmki, Florian Flachsenberg, Patrick Penner, Sophia M. N. Hönig, Tobias Harren, Matthias Rarey

## Abstract

Predicting protein-ligand bioactivities is known to be challenging yet crucial in any drug discovery project. In a protein structure-based scenario, supervised machine-learning models have been highly competitive for at least 30 years. Regardless of the machine-learning method used, dataset size and quality are key aspects in model training and validation. In general, datasets are the foundation upon which accurate performance estimates can be obtained. While well-curated repositories exist for bioactivity and protein structure data, combining these two types of data is particularly challenging. With ActivityFinder, we recently introduced a fully-automated process for linking these data sources relying on protein sequence and molecular structure only. By combining ActivityFinder with previously developed tools for structure quality estimation and property calculation, we created StrAcTable, an automatically constructed dataset of annotated protein-ligand complexes. The automated procedure allows for continued and sustainable growth. StrAcTable includes detailed descriptions of the quality of matching between ChEMBL and PDB, of the macromolecular structure, small-molecule ligands bound, and bioactivity data from ChEMBL. Based on ChEMBL Version 35, the StrAcTable contains 20 063 protein-ligand complexes with bioactivity values, enabling an efficient construction of training and validation datasets for structure-based molecular design method development.

## 1 Introduction

Predicting the bioactivity of a protein-ligand complex is the key challenge in early-phase drug discovery. Computational, structure-based approaches, such as hit identification (virtual screening), target identification (inverse screening), or compound optimization, rely on binding affinity estimates. This includes assessing the drug candidate’s binding affinity to the primary target, but also its selectivity and cause of potential off-target side effects. Despite decades of study,^1–6^ reliable, widely applicable scoring functions are lacking.^7,8^

Multiple issues hinder the progression towards more reliable predictions, yet dataset size and quality are often at least part of the problem. New methods are frequently developed with novel datasets, indicating that no consensus on a gold standard dataset for method development or validation exists.^9–18^ For newly developed datasets, the used data and the methods to collect and combine data from multiple repositories vary between approaches.^12,14,17–19^ Frequently, critiques and opportunities for improvement are expressed on specific datasets^9,20–23^ or on the general methodology of data curation and testing.^23–27^ Concordantly, two independent recent reviews stated that dataset size and quality are significant limiting factors for the performance and generalizability of machine-learning methods.^28,29^ The role of datasets in structure-based drug discovery is two-fold. They are used to both train or develop new methods^12,13,18^ and test their performance,^14,15,30^ for machine-learning as well as for traditional scoring functions. Therefore, the underlying data used to construct these datasets is especially important.

Although data quantity and quality could be improved for all kinds of datasets related to drug discovery, those comprising experimentally determined protein-ligand structures and measured bioactivity data are particularly challenging to construct and maintain. While there are well-curated and continuously updated sources for pure structural data (PDB^31^) and bioactivity data (ChEMBL,^32^ PubChem^33^), combining both is not a simple task. Three influential long-standing sources for the combination of structural data and bioactivity data are PDBbind,^34–37^ BindingMOAD,^38–41^ and BindingDB.^42–45^ PDBbind and BindingMOAD aim to provide affinity data for all suitable entries in the PDB by searching the primary publication of a PDB entry (PDBbind), and beyond (BindingMOAD). BindingDB primarily screens literature, especially US patents, to collect affinity data that it then automatically crosslinked to PDB entries if possible. BindingDB states that crosslinking is performed using exact ligand matches and sequence alignments, either with 85% or 100% sequence identity, but does not provide further details.^46^

All three approaches have in common that they involve laborious manual efforts (PDB- bind even double checks all entries by two independent researchers), which drastically increases the effort to maintain them and can introduce human error.^47^ In the case of PDB-bind, further manual checks are performed to filter towards higher quality levels by checking for sufficient electron density of the ligand, for example. ^36^ Using manual steps to curate a dataset with ever-growing experimental data complicates its perpetuation. BindingMOAD announced that they would cease their efforts in maintaining their database,^47^ and PDB-bind switched to a paid model to continue efforts to keep up to date with the increasing volume of literature data. With these two remarkable and commendable efforts to provide free data to the public ceasing, there is an increasing need for continuously updated data on protein-ligand complex structures with annotated bioactivities.

Two recent developments are the Papyrus Dataset^17^ and BioChemGraph^48^, which both link PDB and ChEMBL using UniProt IDs and InChIKeys. Despite being automatic, these approaches underestimate the amount of data that could be linked between repositories (e.g., by not linking data of racemic mixtures), have issues with the quality of sequence matches using UniProt IDs,^49^ and do not include options to filter out low-quality structures.

This publication presents a new dataset, the Structure Activity Table (StrAcTable). The recently published ActivityFinder^49^ enables automatic crosslinking of structural and bioactivity data and was applied to PDB and ChEMBL. ActivityFinder reads the information on sequences in PDB structures and bioactivity assays, respectively, as well as information on chemical structures found in both resources. Parts of the sequence near the molecule of interest in the PDB structure are handled with special care, recording exact differences. ActivityFinder builds molecules from their 3D-coordinates in PDB files to most closely resemble the modeled data. However, using this approach, common errors in the modeled structure might lower the quality of the resulting dataset.

LigandExtractor, which finds any potential ligands present within a structure, is employed to help offset this problem. The methodology behind LigandExtractor was first descriped in *Flachsenberg et al.*^50^ All potential ligands are checked for compatibility with the chemistry model of the NAOMI cheminformatics library.^51^ Furthermore, any potential problem annotated in the metadata (header) of the PDB file as well as inconsistencies of the interpreted ligand structure with the metadata are reported.

StructureProfiler^52^ assesses the quality of the overall PDB structure. It automates many tests of structural quality, outputs multiple quality criteria for the structure, some ligand descriptors, and calculates the support of the complete structure, the binding site, and any ligands within the electron density. Utilizing StructureProfiler and LigandExtractor, we can automatically assess the quality of a PDB structure and modeled ligands. With these three software tools, we have developed StrAcTable that combines bioactivity data from ChEMBL and structural data from PDB, including information on ligand completeness, type (e.g. organic, covalent), and various quality criteria on structures, ligands, and bioactivities. Quality descriptions are provided for matching the structure and activity data for protein sequences and small molecules. StrAcTable is a new, highly flexible resource for the scientific community, enabling both machine-learning and traditional docking-scoring development. Due to its automation, updates can be provided with substantially reduced manual efforts. Furthermore, proprietary in-house data can be used to enrich and customize StrAcTable.

## 2 Methods

The Structure Activity Table (StrAcTable) is designed as a next-generation dataset for structure-based bioactivity prediction. StrAcTable is derived in a fully-automated fashion from ChEMBL and PDB using a collection of tools from the NAOMI ChemBio Suite. While the primary use of StrAcTable is the creation of training and validation data for structure-based design approaches, some application scenarios have specific requirements we address by three additional variants of StrAcTable (see Table 1).

**Table 1:** Description of all four versions of StrAcTable. They differ only in which data is included from ActivityFinder. All contain the same data from StructureProfiler and LigandExtractor.

| Abbreviation | Name | Characteristic |
| --- | --- | --- |
| StrAcTable | Structure Activity Table | Protein-ligand bioactivity data |
| StrAcTableT | Structure Activity Table target | Protein-target bioactivity data |
| StrAcTableF | Structure Activity Table filtered | Filtered subset of StrAcTable |
| StrAcTableTF | Structure Activity Table target filtered | Filtered subset of StrAcTableT |

In case scientists aim to identify the optimal ChEMBL target for a given PDB structure but there are no recorded bioactivities for a present ligand, or all recorded bioactivities are associated to targets with suboptimal sequence matches we provide StrAcTableT. In StrAcTableT, all found ligands are ignored, so all matching ChEMBL targets are returned, enabling users to query for all bioactivity data associated with a target of interest. Additionally, scientists may want to focus on the most accurate data for a protein-ligand pair (PL-Pair) and therefore ignore data from alternative ChEMBL targets. To address this need and enable users to work with the data quickly while preserving all relevant information, we provide the alternative versions StrAcTableF and StrAcTableTF. Only the best-matching ChEMBL targets, as determined by criteria explicitly developed for this purpose, are retained in both datasets. Matches constructed using SEQADV entries in the PDB structures are discarded during this filtering process because the mutations in the SEQADV entries are not present in the modeled PDB structure, by definition. Additionally, we annotate all versions of StrAcTable with easily understandable quality criteria to enable quick quality-based filtering. Several previously developed tools are applied during the generation of all datasets. The following paragraphs provide important details on how these tools work.

### 2.0.1 LigandExtractor

LigandExtractor is a tool first developed to find all potential ligands in a PDB file and performs basic ligand consistency checks. It was first described (as an unnamed in-house tool) in the context of creating the PDBScan22 dataset.^50^ LigandExtractor allows for an automated assessment if there are any particularities or problems with the respective ligand. This includes annotated peculiarities in the PDB file’s metadata (e.g., missing atoms), covalent linkage of the ligand to the protein, problems with the handling of the ligand in NAOMI (e.g., metal-containing compounds), or deviations of the interpreted ligand structure from the metadata (e.g., different chemical formula). Detailed analysis and annotation of a ligand’s particularities allow users to make an informed but automated decision about how to handle specific ligands. A comprehensive list of all skip reasons which annotate these problems or particularities along with exemplary PDB codes for further illustration can be found in the Supporting Information Section 3. In addition, LigandExtractor calculates molecular weight and the number of heavy atoms for all present ligands.

### 2.0.2 StructureProfiler

StructureProfiler^52^ was published in 2019 and is a tool for automated assessment of X-ray structures of protein-ligand complexes. It allows for assessing global structure quality criteria, similar to those in frequent dataset configurations, and enables users to estimate the local fit of parts of the structure to the underlying electron density using EDIA_m_.^53^ The electron densities are generated from 2fo-fc cif files using GEMMI,^54^ as described in the Supporting Information.

### 2.0.3 ActivityFinder

The recently published ActivityFinder^49^ links bioctivity data, in this case from ChEMBL, to structural data from PDB files, offering two modes. The binding activity-focused mode connects protein-ligand complexes to related target-compound binding data found in ChEMBL assays. Protein sequences used to map into ChEMBL can be limited to those that are part of a binding pocket and, therefore, are guaranteed to be near the ligand. All mutations are tracked for the complete sequence for target matching, and all mutations within the binding site, defined by a sphere of 6.5 Å in diameter around any ligand heavy atom, are reported. Target-specific component sequences and assay-specific variant sequences are handled separately. This strategy captures the number of identical amino acids between the protein sequences in a PDB file and their canonical reference, while also evaluating their identity with the construct used in the binding assay. Please refer to *Ehmki et al.*^49^ for a more detailed description. Data from this mode contribute to StrAcTable and StrAcTableF. The target mode considers only the protein part of a protein-ligand complex. All matching ChEMBL targets are recorded, regardless of binding activity data availability or the location of the matching protein sequence in the complex, allowing users to find the best-matching target for a given protein structure in the PDB irrespective of the ligand and bioactivity information. Data from this mode contribute to StrAcTableT and StrAcTableTF. It is important to note that in the current implementation of ActivityFinder, only ChEMBL targets with a sufficient match to any PDB structure containing an existing ligand are recorded. ActivityFinder only considers protein-ligand complexes with at least one valid ligand. That means, even if an apo structure in the PDB has a matching entry in ChEMBL, the ChEMBL target will not be part of an ActivityDB instance because the apo structure is not included during the creation of an ActivityDB.

### 2.0.4 Bioactivity data filtering

Using BLAST^55^ to map PDB sequences to ChEMBL target sequences yields several possible mappings. While this can be beneficial for users who want to investigate all possible datapoints, it complicates the automated selection of the best possible bioactivity data for any given PL-Pair (StrAcTable) or PDB structure (StrAcTableT). To determine the best-matching target, a filtering procedure was developed and validated using a set of pharmaceutically relevant targets based on the quality of the link (see Supporting information Section 6.1). Using this procedure, we create filtered versions of the aforementioned datasets called StrAcTableF and StrAcTableTF, that enable users to work more easily with the most accurate data.

Relying solely on the percent identity of the match tends to favor alignments that cover only small parts of the query or target rather than more desirable ones. To differentiate multiple sequence matchings, a protein matching score is calculated as shown in Equation 1 using the percent identity of the match and the query coverage (*Cov_match_*_+*query*_) that describes how much of the query PDB sequence is covered in the match. Histograms of all used metrics can be seen in Figure S3.

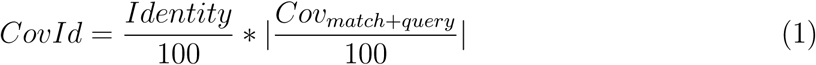

Starting with all possible ChEMBL targets for each PDB structure (StrAcTableT) or PL-Pair (StrAcTable), the following filter cascade is applied for ChEMBL target selection (see Figure 1). First, the Unchecked CHEMBL612545 target is removed as it contains heterogenous data (dummy filter). Second, only ChEMBL targets with the highest *CovId* are kept (sequence matching filter). Third, only ChEMBL targets with the best possible ligand matchings are kept (molecule matching filter). Fourth, the targets with the fewest mutations are chosen (mutations filter). The fifth step utilizes the ChEMBL target type to filter for the most specific ChEMBL targets (target type filter). Targets with the single protein, protein complex or protein family tag are used preferentially in that order. If all targets are annotated differently, this step is skipped. The sixth step retains only those ChEMBL targets with the highest number of associated unique activity values (Datapoints filter). This maximizes the amount of data, as any targets reaching this step in the filter cascade are equally suitable. As a final tiebreaker, the ChEMBL target with the lowest number is chosen to avoid randomness (tiebreak filter). Steps three, four, and six can only be performed if ligand and binding site are known; therefore, they do not affect filtering for StrAcTableTF.

**Figure 1:**
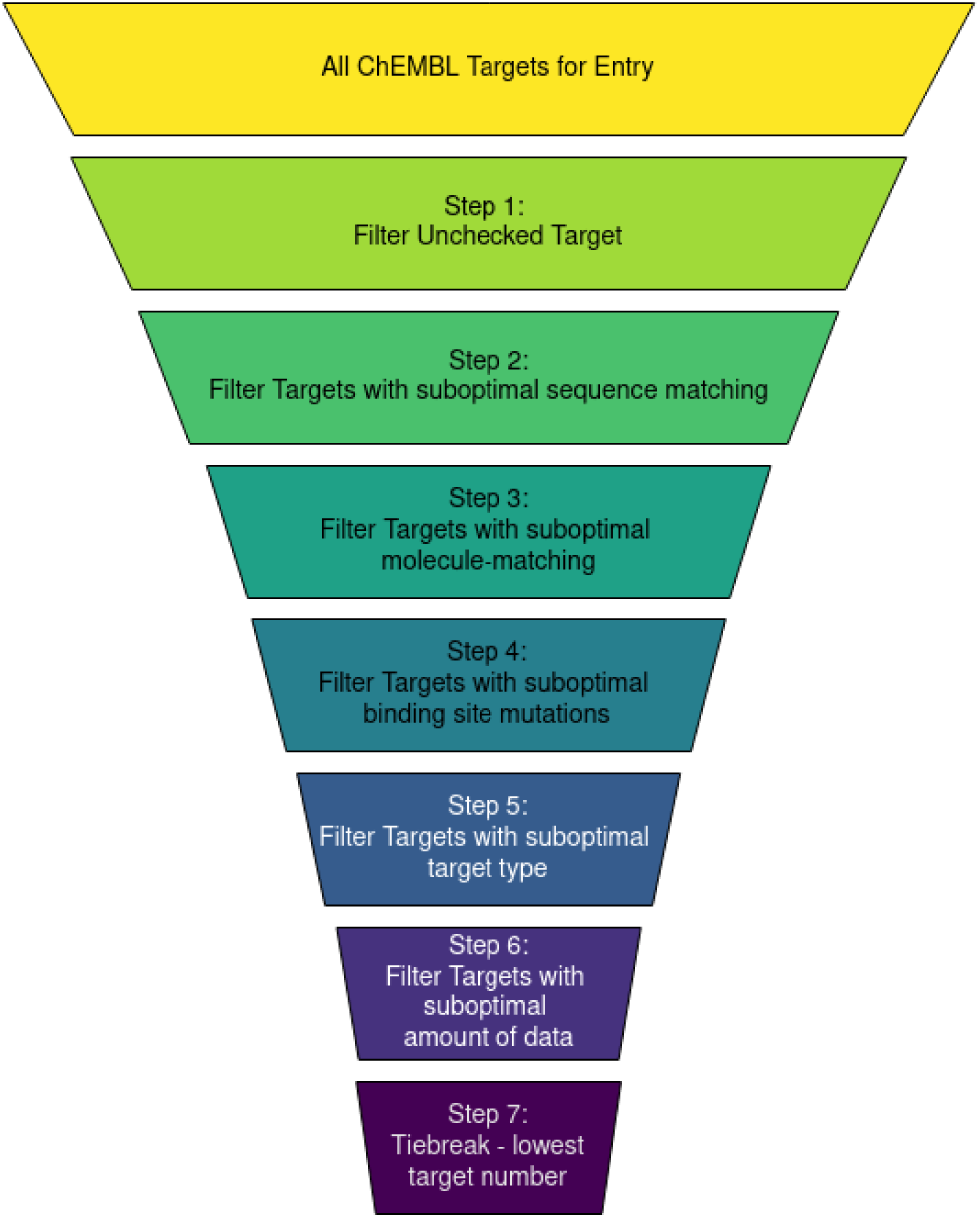
Illustration of the filtering workflow for all ChEMBL targets of a PDB-ligand pair (StrAcTable) or a PDB structure (StrAcTableT). Exact numbers for both filtering procedures are given in Figure 3.

### 2.0.5 Sequence matching quality levels

However, using only the best match does not fully describe its quality, as all suitable ChEMBL targets with a sequence identity of at least 80% are reported. To quickly filter for absolute sequence match quality that goes beyond sequence identity, we developed three categories for use-case-dependent quality assessment (see Table 2). We use the length of the alignment, the used sequence of the PDB structure (query), and either the used component sequence of the ChEMBL target or ChEMBL assay variant sequence (hit), and the percent identity of the match to calculate additional metrics *D_match_*_+*query*_ (see Equation 2) and *D_match_*_+*hit*_ (see Equation 3). Histograms of all used metrics are displayed in Figure S3.

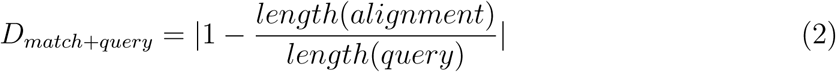

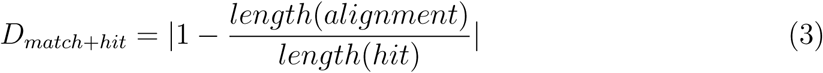

**Table 2:** Quality levels of protein matching

| Group | Sequence Identity | $D_{match+query}$ | $D_{match+target}$ |
| --- | --- | --- | --- |
| Gold | 95% | 5% | 5% |
| Silver | 95% | 10% | - |
| Bronze | 80% | - | - |

The Gold level encompasses all protein matchings in which the PDB and ChEMBL sequence are a precise match and are similar (difference below 5%) in size. This level is designed for users who want to ensure the highest possible quality of the sequence match. The Silver level expands upon this group by allowing a greater discrepancy in the length of the PDB and ChEMBL sequences. While ensuring that most of the PDB sequence is covered (a minimum of 90%), it allows all longer ChEMBL sequences to be matched. Since many sequences in ChEMBL are far longer than those in a classic PDB structure (e.g., the *HIV* polyprotein sequence versus only the protease in a PDB structure), this level enables users to establish these connections when sufficient sequence identity is present. Lastly, the Bronze matching level encompasses every match with at least 80% sequence identity.

### 2.0.6 Workflow of data generation

The workflow for generating the StrAcTable consists of five major steps. First, ActivityFinder, StructureProfiler, and LigandExtractor are executed in parallel for all PDB entries to be investigated. The second step merges the respective output files to a single one for each tool and type of output, which includes the primary output of the respective tools and secondary output of ActivityFinder with the recorded binding site mutation data. In the third step, the bioactivity data is enriched and filtered. The ChEMBL database is queried for additional data for processing (e.g. variant sequence assays or target classification data) or direct addition (e.g. target hierarchy or ChEMBL release information) to the parsed activity data for future analysis. Then, the quality level and score are calculated and added to each entry in the activity data. An additional filtered version of the activity data is generated where only the best-matching ChEMBL target is used for each PDB structure. In the fourth step, the structure data, ligand data, and the two versions of the activity data are combined to create the final StrAcTable and StrAcTableF datasets. One dataset contains all possible ChEMBL targets per PDB (StrAcTable), and the other contains only the best matches according to our developed metrics (StrAcTableF). Similarly, the target mode activity data is used to create the StrAcTableT and StrAcTableTF. In the final step, the number of mutations in the binding site as annotated by ActivityFinder is added to StrAcTable and StrAcTableF, regardless of mutation type. If a variant sequence exists for an assay, the match to the component sequence is ignored in further analysis for that assay. Consequently, if the variant sequence includes a mutation consistent with a mutation in the PDB structure sequence, no mutation is recorded. In ChEMBL Version 35, there are 17070 assays with a variant sequence, 1861 (10.90 %) of which have the variant sequence UNDEFINED MUTATION. As no usable sequence is annotated to the variant sequence UNDEFINED MUTATION, all assays annotated with this mutation are handled like assays without a variant sequence annotation. An additional filtered version of the bioactivity data is generated where only the best-matching ChEMBL target is used for each PDB structure (StrAcTableT) or PL-Pair (StrAcTable). As long as there are no changes to the schema of the ChEMBL or changes in the PDB format that prevent NAOMI^51,56^ tools from reading them, this process can be used for automated regular updates.

## 3 Results

Four different versions of StrAcTable have been developed in this publication (see Table 1). The following Section examines the data present in all versions, the developed target filtering metrics, and showcases how to use StrAcTable.

### 3.1 General information about StrAcTable

There are 20 063 protein-ligand complexes for which we can annotate an activity value. 13 042 of 51 678 unique molecules have at least one activity value associated in the StrAcTable. StrAcTable, prior to any filtering, comprises 3 619 313 rows and 134 columns while being a full outer join of the data of LigandExtractor (12 columns), ActivityFinder (85 columns), and StructureProfiler (36 columns), two merge indicators, and additional quality columns. A detailed list of all columns in StrAcTable, including a description, is given in the Supporting Information. The target-centered versions StrAcTableT and StrAcTableTF have a reduced number of ActivityFinder columns (37) and therefore only 85 total columns. For the creation of StrAcTable, ChEMBL Version 35 and a PDB mirror from the 04/25/2025 were used, containing 226339 total PDB entries. Table 3 shows statistics for all StrAcTable versions. This includes the number of PDB entries with matched ChEMBL data, the number of unique molecules, and the number of complexes with and without activity data.

**Table 3:** Statistics on the number of rows, columns, PDB entries, unique molecules and unique Complexes. All columns with an * only count entries where ChEMBL data was available.

| Descriptor | StrAcTable | StrAcTableT | StrAcTableF | StrAcTableTF |
| --- | --- | --- | --- | --- |
| Rows | 3 619 313 | 27 113 695 | 2 271 838 | 2 847 075 |
| Columns | 134 | 85 | 134 | 85 |
| PDBs* | 19 262 | 48 296 | 19 237 | 48 206 |
| Molecules | 51 678 | - | 51 678 | - |
| Molecules* | 13 042 | - | 13 037 | - |
| PL-Pair | 557 853 | - | 557 853 | - |
| PL-Pair * | 20 063 | - | 20 033 | - |

StrAcTable records multiple measurements for each PL-Pair if they are present in ChEMBL. As shown in Figure 2a, the vast majority exhibit one or two distinct recorded activities. However, there are some PL-Pairs with more than 900 different measurements, for example, the drug Vorinostat in PDB 4LXZ.^57,58^ While in the majority of cases, each PDB only has a single ChEMBL target associated with it, in some cases, there can be up to 20 different ChEMBL targets with sufficient sequence identity (see Figure 2b).

**Figure 2:**
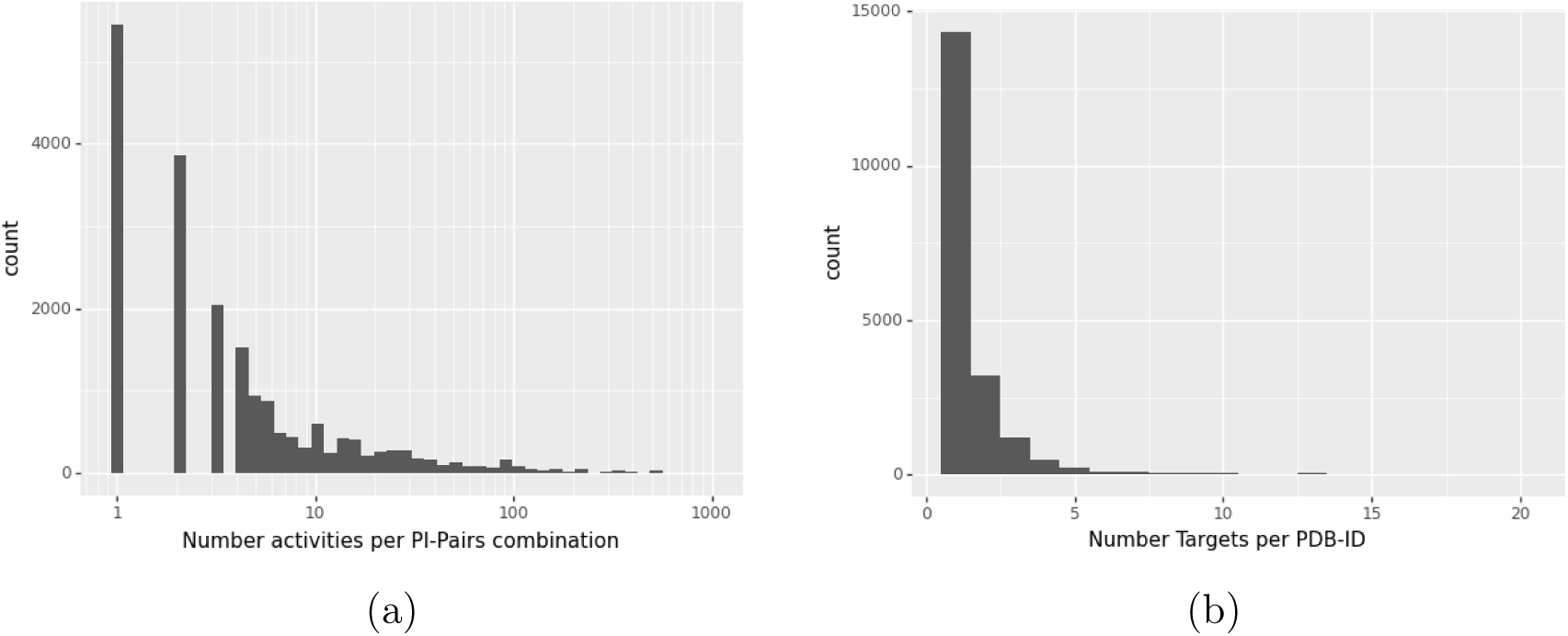
Number of occurrences of PDB-ligand complexes with a specific number of different activities (logarithmic scale) (a) and number of different ChEMBL targets per PDB (b) in the activity data.

### 3.2 Complete StrAcTable filtering statistics

For many PDB structures, more than one possibly matching ChEMBL target is found, and sometimes, multiple different BLAST matches are available for each possible ChEMBL target. For those users only interested in the best-matching data and therefore ChEMBL target, StrAcTableF and StrAcTableTF are provided. A detailed description of the filtering cascade is provided in Section 2.0.4 and detailed results on a set of pharmaceutically relevant targets is given in SI Section 6.1.

The statistics of the filtering cascade are shown in Figure 3. Afterwards, only the best possible BLAST match is allowed for each complex. The exact numbers are provided in Supporting Information. For StrAcTable, 20 033 PL-Pair combinations with suitable activity could be matched, and in 14 647 of these cases, only a single target is found. The small difference in the number of PL-Pairs compared to Table 3 is due to PL-Pairs that only match using SEQADV sequences which are discarded in filtering. For the remaining 5 386 combinations, the filtering pipeline is applied. Combinations are decided mostly using protein matching quality and, to a lesser extent, protein target hierarchy. The target matching tiebreak is only used in 36 combinations across the entire PDB.

**Figure 3:**
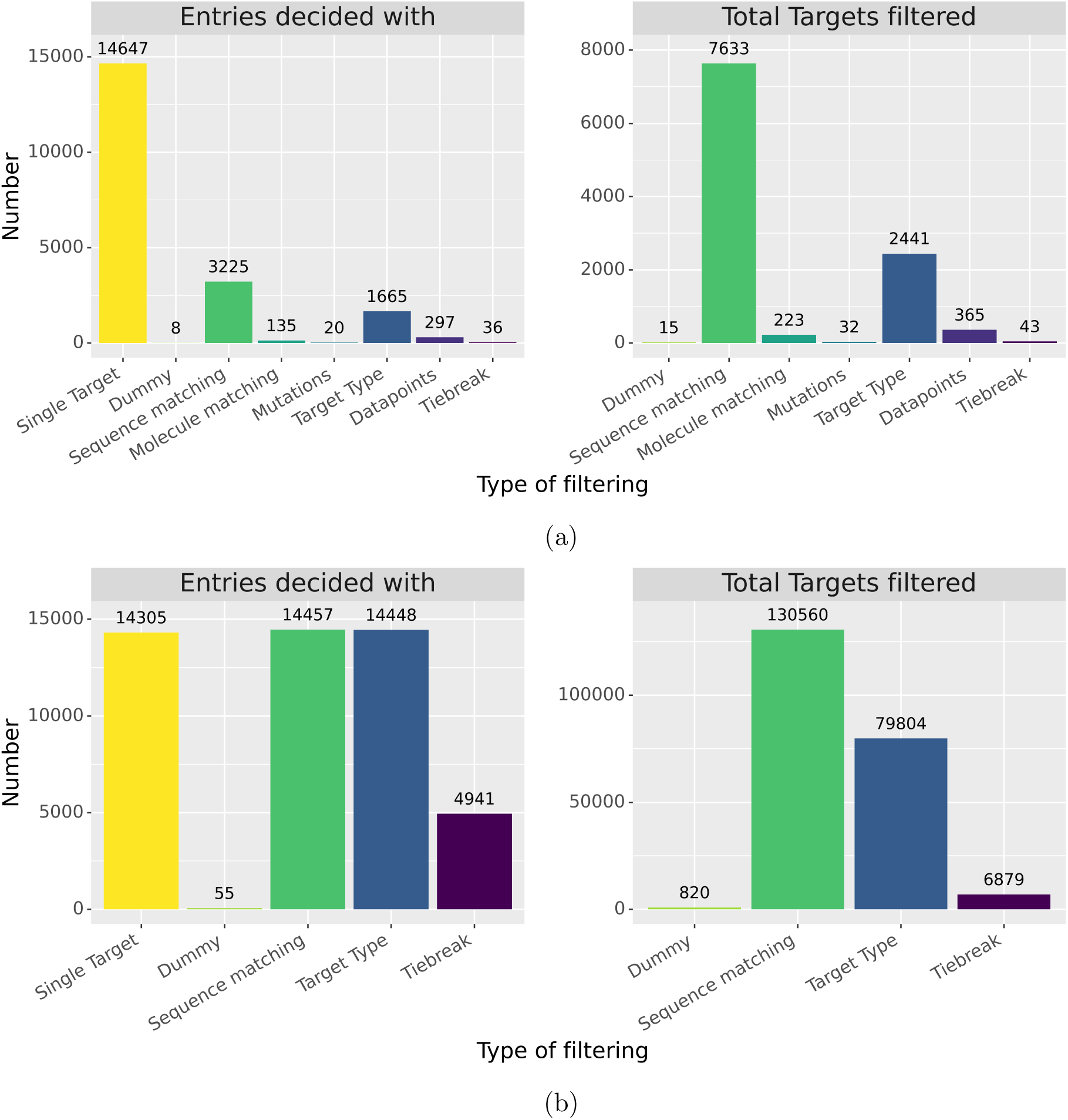
Filtering data for a) StrAcTable and b) StrAcTableT. The left side of each graph shows all PL-Pair (a)/PDBs (b), indicating which step in the filtering cascade the final target could be assigned. The right side of each graph shows the number of ChEMBL targets discarded by each step of the filtering cascade. Dummy describes the Unchecked target as described in Section 2.0.4. that our filtering process removes surprisingly many bioactivity data points when filtering them down to a single target.

For StrAcTableT, there are many more PDB entries for which we can identify possible ChEMBL targets (48 206), but only a small subset (14 305) has a single ChEMBL target. This makes sense, as the number of potential ChEMBL targets is much larger if not restricted by a measured bioactivity of a specific ligand. A roughly equal number of complexes are determined by protein matching quality (14 457) and ChEMBL target hierarchy decisions (14 448). In addition, for many more targets (4 941), a tiebreak is necessary for a final ChEMBL target to be assigned. This is due to the increased number of targets found and the reduced number of possible elements in the filtering cascade.

### 3.3 Composition of the StrAcTable

All versions of StrAcTable are full outer joins of the data from LigandExtractor, Structure-Profiler, and ActivityFinder. Therefore, many entries lack data from certain tools, such as when no matching bioactivity data is found in ChEMBL. Venn diagrams of the contributions of each tool to StrAcTable and StrAcTableF are shown in Figure 4. Data from all three tools are present for only 12.44 % of the entries in StrAcTableF. Contrary to that, the unfiltered StrAcTable contains 38.32 % entries in which all three tools contribute data. This means

**Figure 4:**
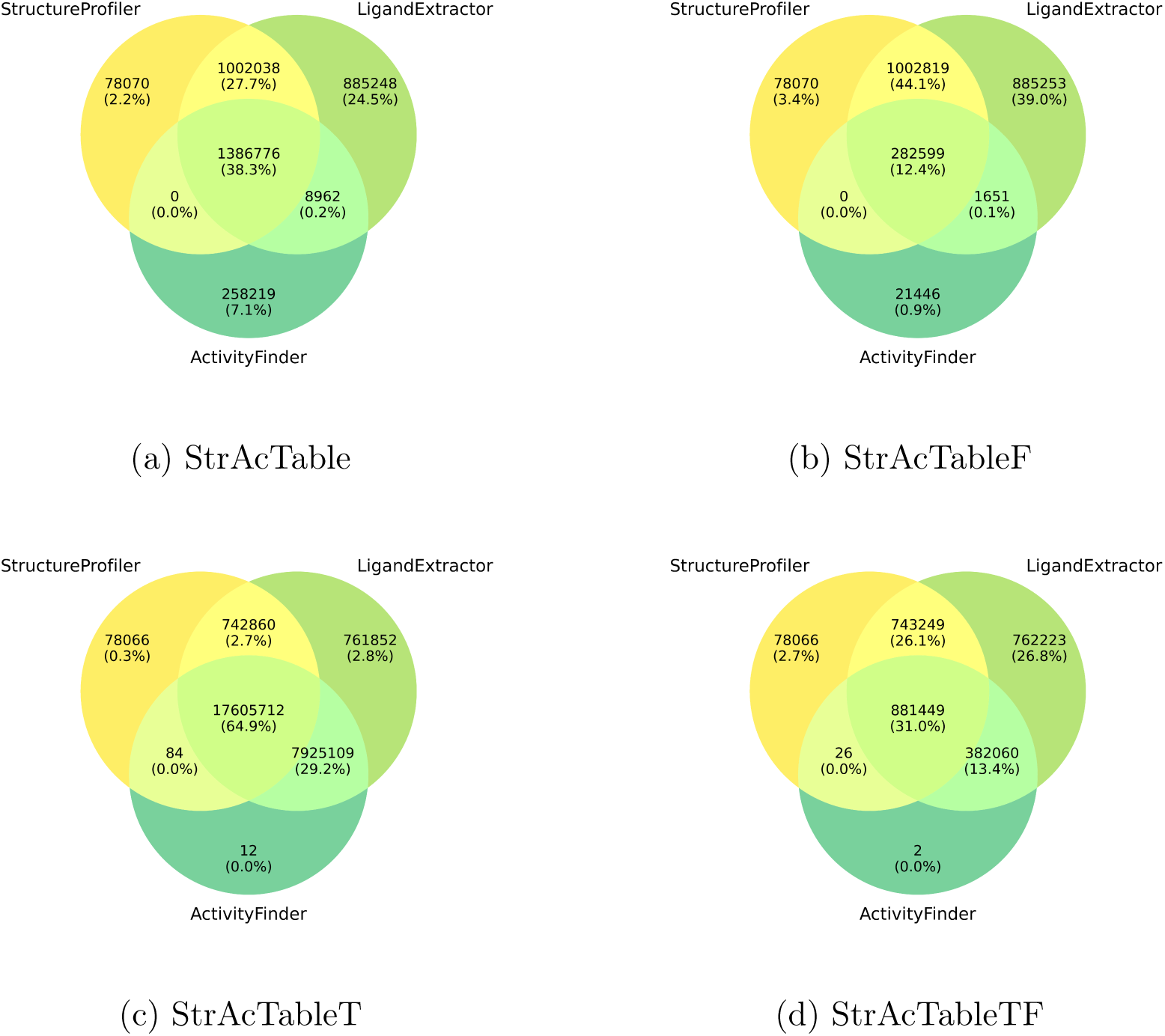
Venn diagrams of StrAcTable statistics, which tools contribute which percentage of entries to it for StrAcTable (a), StrAcTableF (b), StrAcTableT (c) and StrAcTableTF (d).

The following analysis is solely done for StrAcTableF. For most application scenarios, StrAcTableF is the most reasonable dataset since it is not inflated by activities of the same molecules in different ChEMBL targets. For StrAcTableF, most entries either comprise data from LigandExtractor and StructureProfiler or solely from LigandExtractor. This is intuitive, as most ligands (e.g., crystal additives) found in PDB will not have any recorded activity in ChEMBL but still have information on the structure quality and ligands present. The high number of entries with data only coming from LigandExtractor is a result of LigandExtractor being designed to record any ligands present in the structure, even if StructureProfiler or ActivityFinder cannot handle them due to reasons like unsupported valences by the NAOMI chemistry model. The remaining 3.44 % entries with data from Structure-Profiler only result from structures without any bound ligands and very few differences of names for polymeric ligands. Note that there are only EDIA_m_ values and related tests for structures for which an electron density is available, even if other data from StructureProfiler is present. For 0.94 % entries, only activity data is present. This is due to alternative binding chains found by ActivityFinder not found by other tools. Both LigandExtractor and StructureProfiler only annotate a ligands native chain while ActivityFinder can find matches to additional close chains, resulting in the alternative binding chains.

### 3.4 Exploring common properties in StrAcTable

To enable automated quality estimation of activities, structures, and ligands, many related descriptors are either calculated or extracted based on the data in PDB and ChEMBL. Visualizations of the distribution of commonly investigated properties, such as the number ofrotatable bonds^59^ and molecular weight of all unique PL-Pair entries are displayed in Figure 5, including a pie chart of the frequency of organisms. A further analysis of all PDB-HET entries is displayed in Figure 6.

**Figure 5:**
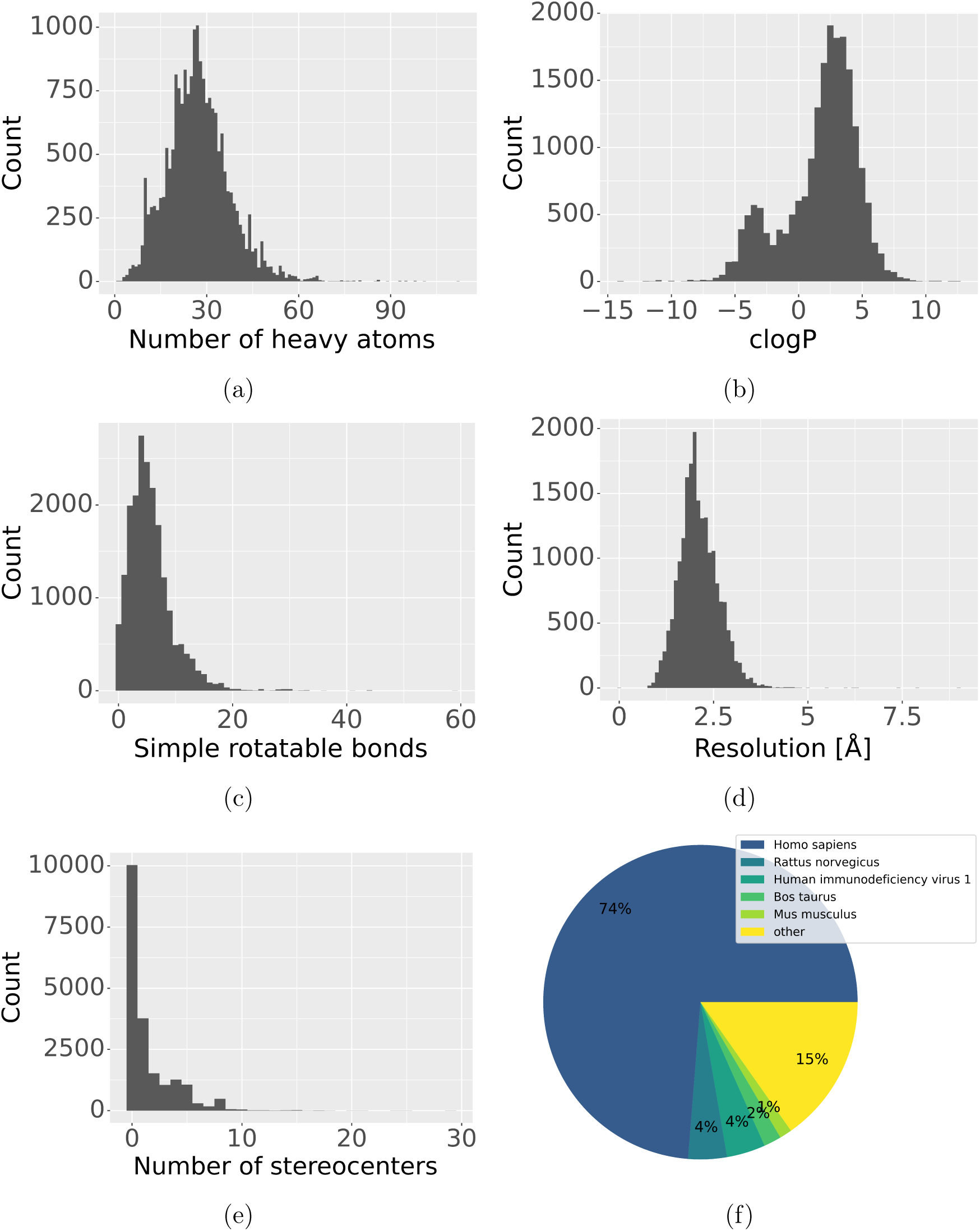
Analysis of the occurrences of the number of heavy atoms (a), calculated logP after Wildman and Crippen^60^ (b), number of rotatable bonds^59^ (c), resolution of the structure (d), number of stereocenters (e), and organisms (f) for all 20 033 unique PL-Pairs in StrAcTableF containing data from all respective tools.

**Figure 6:**
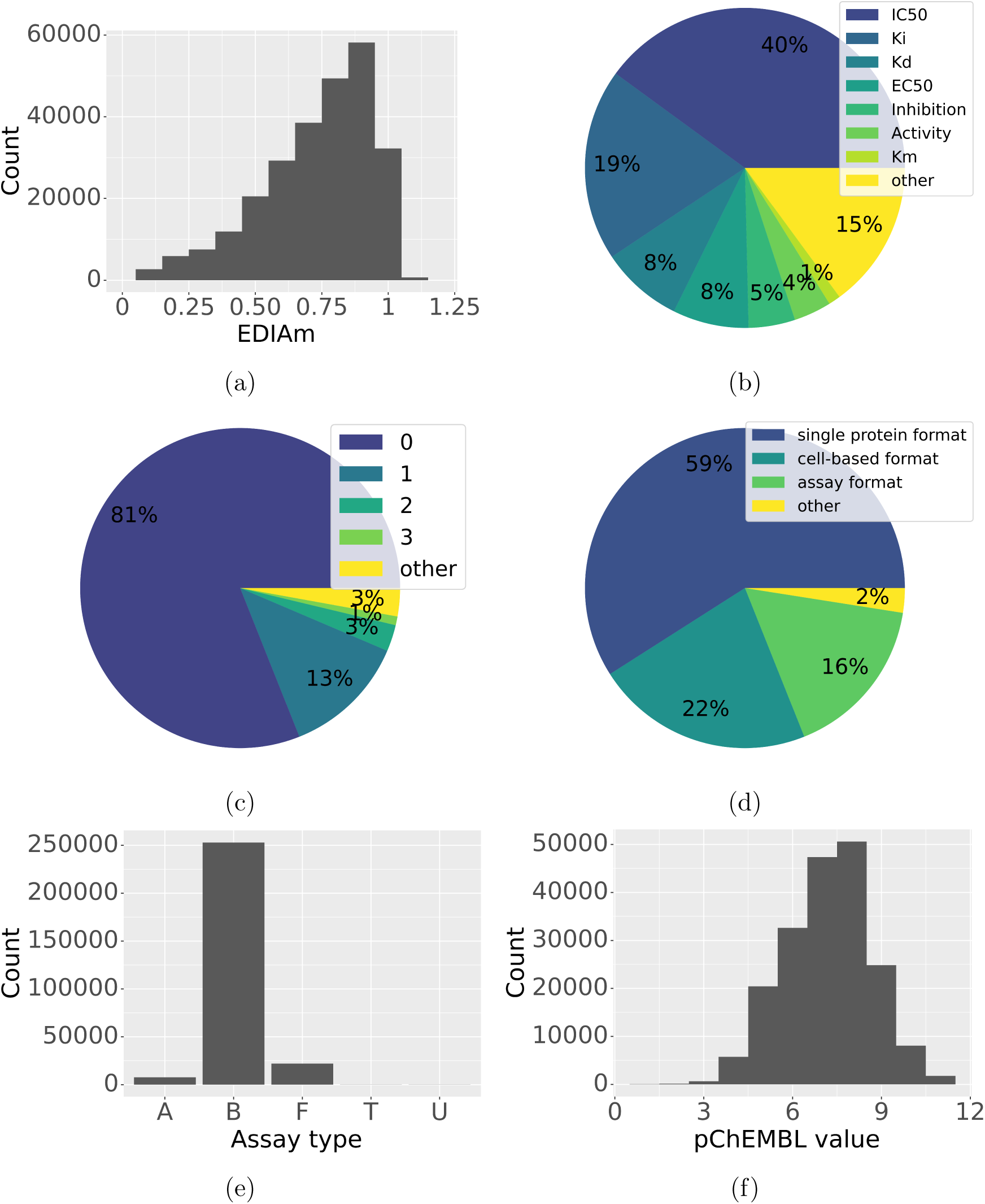
Analysis of the occurrences of the EDIA_m_ (a), the activity types (b), number of mutations (c), Bioassay ontology format^61^ (d), assay types (e), and pChEMBL values (f) in StrAcTableF containing data from all respective tools. Assay types include A (ADME), B (Binding), F (Functional), P (Physicochemical), T (Toxicity), and U (Unassigned).

As shown in Figure 5, the compounds covering 13 037 unique HET codes and 20 033 PDB-HET code complexes exhibit a broad distribution for the number of heavy atoms, clogP after Wildman and Crippen,^60^ number of rotatable bonds, and the number of stereocenters, indicating a high chemical diversity within the dataset. The most common organism found in StrAcTableF is *Homo sapiens*, followed distantly by *Rattus norvegicus*, *HIV-1* and *Bos taurus*. When not correcting for unique PDB-HET code complexes, targets with many activities in the ChEMBL like *HIV-1* have a much higher frequency (see Figure S1e).

Activity types, as shown in Figure 6b, are heavily dominated by *IC*_50_ and *K_i_*values, but very diverse activity types are present in StrAcTableF. As seen in Figure 6a, 118 004 complexes have a high quality EDIA_m_ between 0.8 and 1.2, with declining numbers of complexes showing lower EDIA_m_ and few showing higher values. There are 3 377 complexes for which StructureProfiler can not correctly calculate an EDIA_m_ and returns −1 as a value, which are omitted in the visualization for clarity. 58.54 % of entries in StrAcTableF have been measured in a single protein assay format (Figure 6d), with the rest either being measured in the unspecific assay format or a cell-based format, and 97.95 % have been measured against a target of the single protein target type (Figure S1d). 77.23 % of entries have been curated through autocuration (Figure S1a) and 95.08 % are extracted from a publication (Figure S1b). Assay types are heavily dominated by binding assays, with significantly fewer functional assays, ADME assays, and minimal other types (Figure 6e). As seen in Figure 6f, a wide variety of activity values (mean 7.19, median 7.28, STD 1.45) can be found in the StrAcTable, with both high and low-affinity measurements present. Using ActivityFinder provides end-to-end mutation tracking, as described in the methods section. Mutations in the ligand-binding site are recorded, and their number is annotated to each entry in StrAcTable and StrAcTableF. Notably, the majority of entries (80.64 %) show no mutations in the binding site (see Figure 6c), and 96.67 % have three or fewer mutations.

### 3.5 ChEMBL target distributions

In Figure 7, the first two hierarchical levels in StrAcTableF of the ChEMBL target are shown. An interactive version enabling the visualization of user-filtered data and similar plots can be found in the Supporting Information. Additionally, in Table 4, the exact numbers for the first level of hierarchy are given.

**Figure 7:**
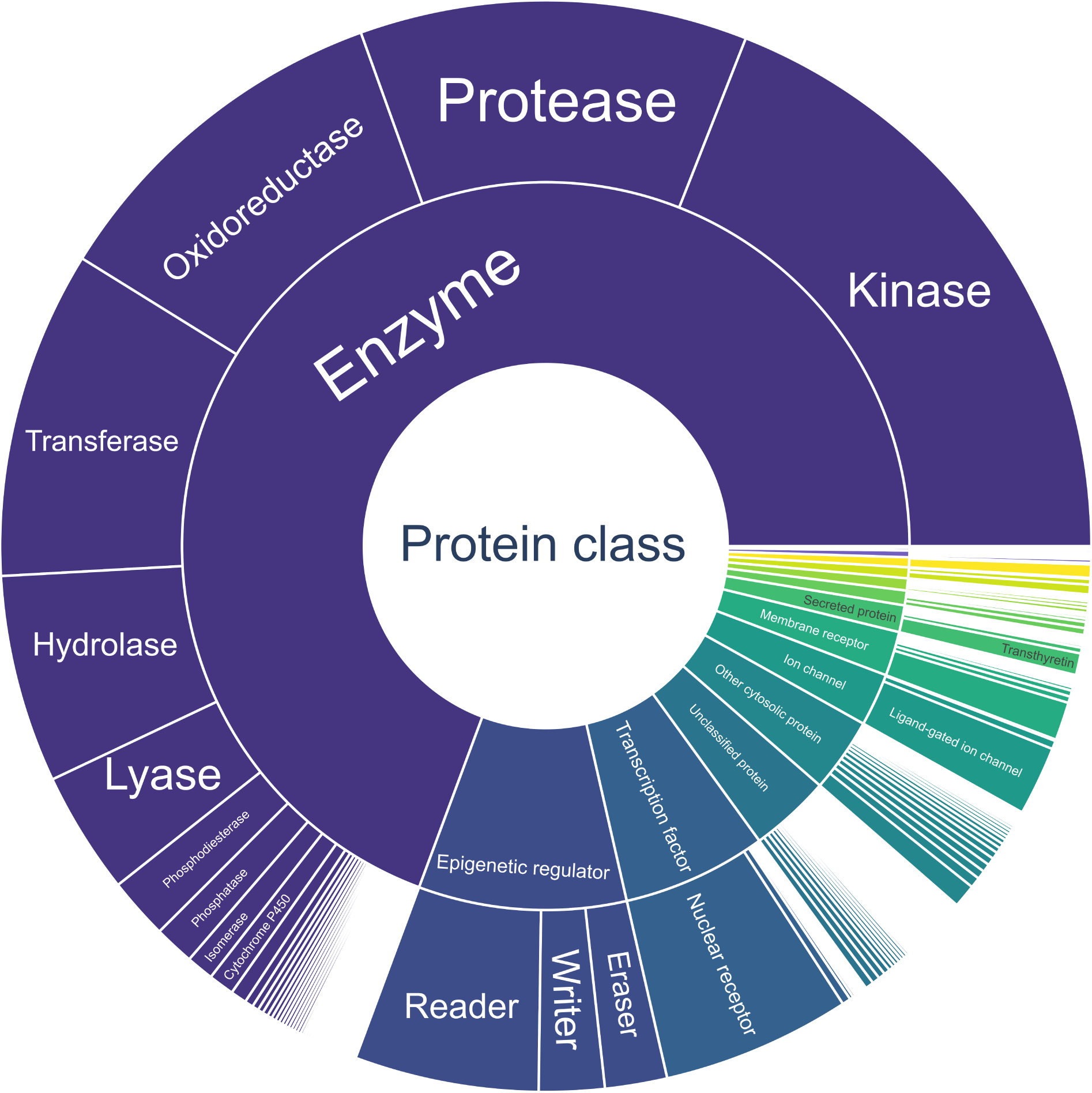
Sunburst plot of the target hierarchy of the ChEMBL targets of all PDB structures in StrAcTableF. For visual reasons, only the first two hierarchical levels are displayed, and any labels that do not fit their boxes are hidden. Complete interactive versions of this figure and any raw data are given in the Supporting Data.

**Table 4:**
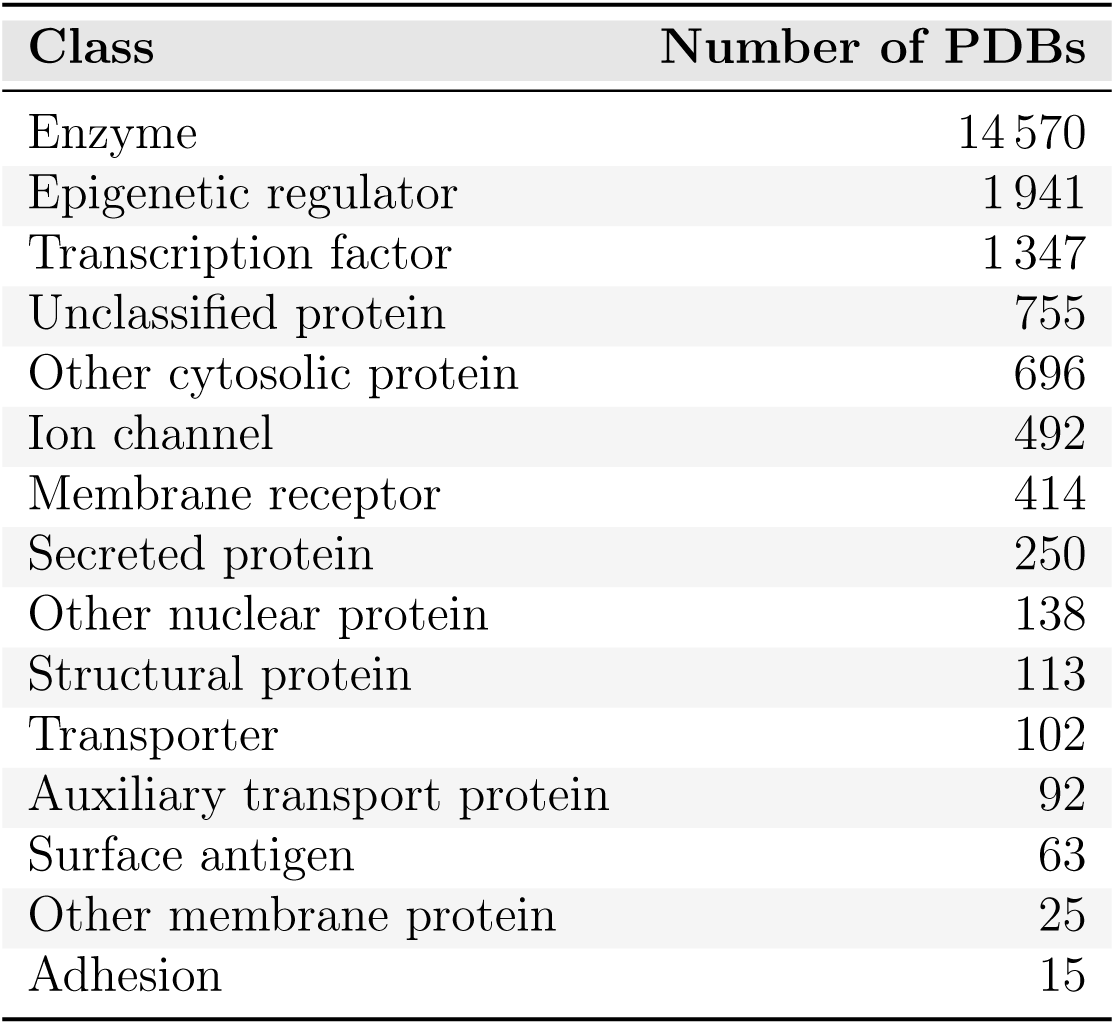
Number of PDB structures found for each first level ChEMBL hierarchy class for the StrAcTableF.

Analyzing Figure 7 and Table 4 reveals that the dataset is highly diverse; it includes data on ion channels, epigenetic regulators, transcription factors, membrane proteins, and many other target classes. However, it is similarly clear that common biases found in PDB and ChEMBL are repeated here, with 19 % of structures categorized as Kinase. Other commonly explored targets, such as Thrombin and Carbonic anhydrase 2, are also frequently found.

### 3.6 Analysis of molecule and protein matching quality

We developed sequence matching quality levels for the PDB to ChEMBL target matching to allow intuitive quality-based data subselections. The protein matching levels are Gold, Silver, and Bronze, with the exact definitions provided in the Methods section. Ehmki *et al.*^49^ introduced five molecule matching levels to categorize matches between PDB and ChEMBL. They can be generally classified into three groups: identical molecules (identical InChI key / 5 and USMILES with chirality / 4), stereochemically distinct molecules (canonical USMILES lacking chirality / 3), and potentially identical molecules (clipped InChI atom and connection layers / 2, or clipped InChI atom layers / 1).

As can be seen in Figure 8d, 86.98 % of the sequence quality in StrAcTableF falls either into the Gold (33.09 %) or Silver (53.88 %) category while only 58.13 % of StrAcTable fall into the same categories. The filtering process enriches high-quality links, even though the quality levels are not directly used. In 22.15 % of the cases, it is possible to find a ChEMBL target with the highest possible sequence quality level and a perfectly matching ligand in StrAcTableF. There is a high frequency of Silver matches since, in many cases, only ChEMBL targets with a more extended sequence than the one modeled in the PDB structure are identified. For ligand matchings, the vast majority of entries match perfectly or almost perfectly (Levels 4 and 5) between ChEMBL and PDB in StrAcTableF (65.77 %) and in StrAcTable (68.06 %), while another 16.90 %/15.66 % match when neglecting chiral information (different isomer) and 17.33 %/16.27 % match with more severe differences.

**Figure 8:**
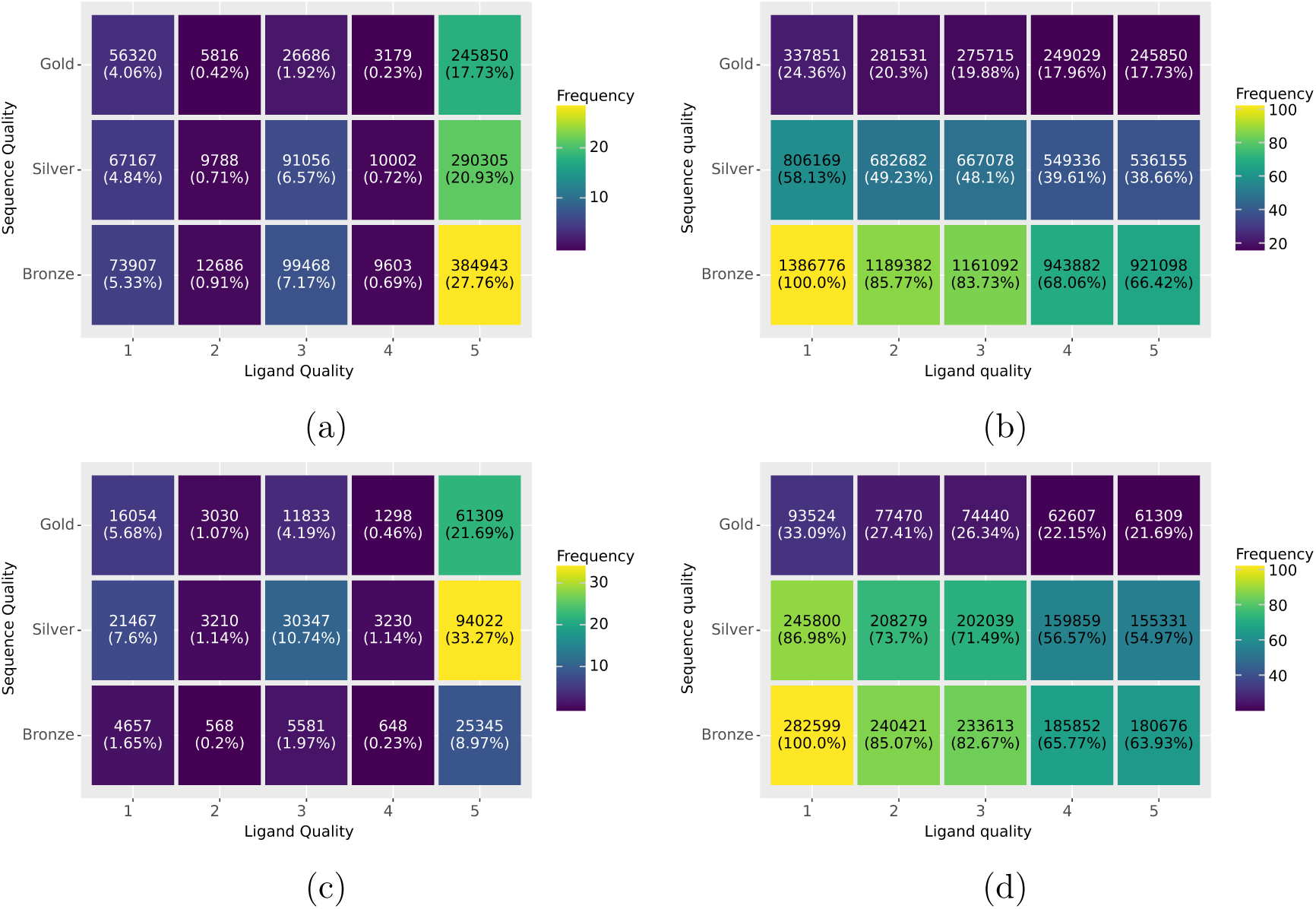
Frequency of protein and ligand matching quality levels in dependence to each other in StrAcTable (a,b) and StrAcTableF (c,d). The exact percentage for any given combination are given in (a) and (c) and the cumulative percentage of the given combination and all qualitatively stricter combinations are given in (b) and (d). Qualitatively stricter combinations mean that both sequence and ligand matching quality are simultaneously higher or equal. Ligand quality levels are 1) truncated standard InChI matches if cut after atom connection layer 2) truncated standard InChI matches if cut after hydrogen connection layer 3) canonical smiles without chiral information annotated 4) canonical smiles with chiral information annotated and 5) standard InChI-key matches.

### 3.7 Growth of the StrAcTable

The process of StrAcTable construction is designed to be as easily expandable to new data points as possible. To showcase how StrAcTable would have grown in the past, we can simulate different past releases of ChEMBL and PDB. Using the later date of the submission of the PDB structure and the submission of the document that was then recorded to ChEMBL, it is possible to estimate how the number of unique PDB-ligand complexes with recorded data in the ChEMBL would have changed over time in a hypothetical scenario where all ChEMBL data is released immediately. As seen in Figure 9, the yearly and cumulative data start to grow significantly in 2005. By 2010, the number of protein-ligand complexes with at least one recorded entry in the ChEMBL grows linearly. The dip in recent years is probably because the used ChEMBL release was in December 2024, and bioactivity and structure data may not have been uploaded or recorded at the same time.

**Figure 9:**
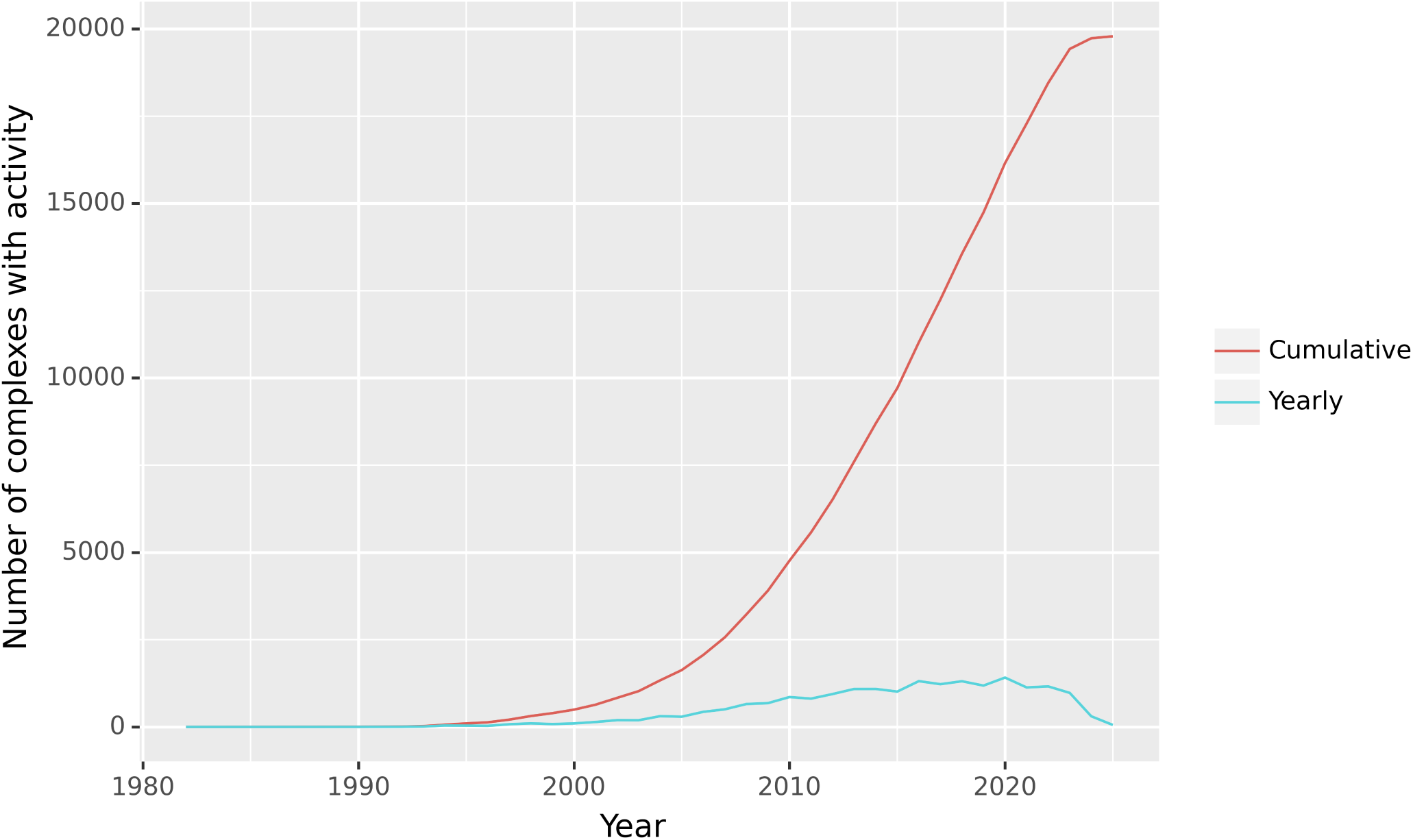
Number of complexes with at least one activity in StrAcTableF for each year. A datapoint is recorded as soon as both the PDB structure and the document out of which the ChEMBL data was extracted was first published. Both are annotated on a yearly basis and cumulatively.

### 3.8 Using StrAcTable for dataset construction

StrAcTable is designed to create novel datasets. To showcase how to collect and use data from the different versions of StrAcTable we explore a single ChEMBL target as an application scenario. CHEMBL1862 is the Tyrosine protein kinase ABL in the single protein human form in ChEMBL. Filtering down StrAcTableF to our target, we find 692 rows. Investigating this further, there are 37 unique PDBs, 42 molecules, 139 ChEMBL assays, and 293 combinations of all aforementioned identifiers. One option is to filter the existing dataset down to a highquality dataset. Several factors should be considered in this case. The first concerns are about the integrity of the structure and the modeled ligand. To account for the integrity of the ligand, we can use the LigandExtractors skip reasons (see Supporting Information Section 3). To account for embedding in electron density, we can use EDIA_m_. However, as EDIA_m_ only works on crystal structures, using this approach, we filter out any structures that were resolved using different methods. These steps filtered out 42 datapoints from StrAcTableF, leaving us with 650 rows. Next, we can filter for global structure quality, for example, the resolution to be below 2.5 Å, after which we are left with 524 rows. The next step is to filter the quality of bioactivity data, removing any potential duplicates and records with data validity comments, after which we are left with 425 rows. Next, we need to consider the quality of mapping the structure to bioactivity data, which again consists of sequence and molecule mapping. For molecule mapping, the best practice is to filter to cases with identical molecules, leaving us with 352 rows. Lastly, there is sequence matching, with two essential criteria: percent identity and the number of mutations in the binding site. One could argue that both are needed, but there is also a valid argument for considering only binding site mutations. If mutations are far outside the binding pocket, they might be less likely to influence binding. When filtering for a sequence identity of over 95% and zero mutations, 337 rows are remaining. Interestingly, we filter out 15 cases with mutations and zero with sequence identity. This highlights again the importance of accounting for mutations in the binding site.

Although the main application of StrAcTable is the construction of large datasets for docking and scoring method development, it can be used for more individual analyses as well. We can focus on a single PDB structure, for example, the T315I mutant structure with PDB-ID 3QRJ.^62,63^ In this case, we only find data for a single assay in our filtered highquality data, and two datapoints with the same activity value, differing only in the exact ligand matched in the PDB file. Notably, the assay contains a variant sequence; therefore, that sequence was matched. If we now look into the unfiltered table, we find 26 extra entries for 3QRJ and five assays compared to the one in the filtered version. This is because we filter not only to the best ChEMBL target, but also to the best BLAST match for the filtered version. Only the assay CHEMBL5108948 aligns best with the mutated structure 3QRJ, as the variant sequence contains the mutation.

An advanced possibility is to investigate if we find good additional data in StrAcTable. Investigating the differences to the filtered version, we find 3 519 extra rows in StrAcTable. We find four new PDB structures in StrAcTable, which means there are better ChEMBL targets for each of these structures. Investigating only the data of these new PDB structures, we can see that the minimum and maximum sequence identity is 84.39 %/99.30 %. The PDB entry responsible for the 99.30 % sequence identity is 3K5V,^64,65^ which is the *Mus musculus* version of Tyrosine protein kinase ABL and therefore matched to CHEMBL3099 with an even higher sequence identity.

Another advanced option is to explore whether we can find interesting apo structures or structures with ligands that lack activity in StrAcTableT by quering target data only. We find 5 967 cases and with that find 18 new PDB entries for that specific ChEMBL target. Examples of this include PDB structures like 8I7T,^66,67^ where there is a binding ligand for which we do not find bioactivity data, or 2G2I,^68,69^ which is an inactive structure with only ADP bound.

## 4 Discussion

Creating an automated workflow to crosslink structural and bioactivity databases is integral for the future development of any method predicting protein-ligand bioactivities. Existing manual approaches for solving this problem have been the backbone of method development for decades and will continue to be critically important. Despite that, solutions need to be able to grow with the continually increasing growth of structure and affinity data generation. Within this work, we automatically generated such a dataset and thoroughly analyzed the present data.

There are some valid opportunities to improve ActivityFinder, StructureProfiler and LigandExtractor and therefore the StrAcTable workflow. As ActivityFinder currently focuses solely on X-ray structures, bioactivity values of cryo-EM or NMR structures are neglected. The EDIA_m_ requires an electron density calculation and is used as a descriptor for the experimental support of the used structure. Therefore, structures without electron density, such as those created using NMR spectroscopy, are discarded when filtering with EDIA_m_, thereby eliminating potentially valuable information. For cryo-EM maps, the Q-Score^70^ was derived from EDIA_m_ and is planned to be included in the future.

So far, NAOMI^51,56^ does not fully process metal-containing ligands. Therefore, any bioactivities of ligands that contain metals are only using a subset of the original ligand, but these cases can be filtered out using the skip reasons. While methods for extracting covalently binding molecules exist, we decided not to take them into account for bioactivity linking in this initial version of StrAcTable, as further problems with matching them to ChEMBL molecules and interpreting the bioactivity need to be addressed. Similarly, enhanced stereochemistry is also not supported so far.

PDB-redo^71^ aims at improving the model quality of PDB structures with a more sophisticated refinement procedure. Creating an alternative version of StrAcTable based on PDB-redo is currently being investigated but is not included in this release.

In addition, for now, only the ChEMBL database has been crosslinked with the PDB, but other databases like PubChem,^33^ BindingDB^42^ would also be valuable additions. As no automatic crosslink between PDB and PubChem exists, and it encompasses both BindingDB and ChEMBL, this would significantly increase the amount and diversity of data available in StrAcTable. While BindingDB crosslinks to PDB entries, the crosslinking is less detailed than the approach used by ActivityFinder. As BindingDB primarily searches US Patents, its bioactivity data is mostly orthogonal to ChEMBL and would further enrich StrAcTable.

## 5 Conclusion

In this work, we present automated workflows for creating datasets with combined bioactivity and structural data. Since the bioactivity of ligands to targets is sensitive to even small changes in the structure, special care was taken to ensure that the target and the small molecule are reasonably similar in both data sources. Any potential differences are reported enabling users to decide on the acceptability of small variations. Furthermore, the experimental evidence for the complex structure was carefully validated.

StrAcTable aims to provide any information a user needs to accurately estimate the experimental support of the structure, activity data, and the matching between the two, allowing for the automatic construction of derived datasets. Due to automation, it is possible to achieve sustainable growth in conjunction with PDB and ChEMBL, eliminating the need for laborious human efforts. The resulting data collection has the potential to be used for improved machine-learning based docking and scoring approaches as well as validation scenarios.

## 6 Data Availability

All Data is availabe through the FDR of the University of Hamburg at https://www.fdr.uni-hamburg.de/record/18244.

## 7 Code Availability

All code used to construct and analyse the StrAcTable, to create the plots in this publication and a jupyter notebook showcasing how to use StrAcTable is available at https://github.com/rareylab/StrAcTable. The LigandExtractor, StructureProfiler and ActivityFinder tool are part of the NAOMI ChemBio Suite which is available at https://uhh.de/naomi, free for academic use and evaluation purposes.

## 8 Author Contributions

Conceptualization: T.G., E.S.R.E., F.F., P.P., S.M.N.H., M.R.; Analysis: T.G., T.H., M.R.; Software development: T.G., E.S.R.E., F.F., P.P., S.M.N.H.; Writing of a first Draft: T.G.; Supervision: M.R. Review and Editing: All authors.

## 9 Acknowledgments

We would like to thank Dr. Christiane Ehrt and Thorben Schulze for valuable discussions about estimating electron density support. We would like to thank Dr. Christiane Ehrt and Paula Kammer for their useful feedback on the use of StrAcTable.

