## Supplementary Information for "Enabling automatic generation of protein-ligand complex datasets with atomistic detail"

‡*Current Address: Pfizer Pharma GmbH, Friedrichstraße 110, 10117 Berlin*

¶*Current Address: BioSolveIT GmbH, An der Ziegelei 79, 53757 Sankt Augustin, Germany*

§*Current Address: Global Discovery Chemistry, Novartis Biomedical Research, Basel, Switzerland*

#### 1 StrAcTable Documentation

An exact documentation on each column in the StrAcTable and therefore each column in the respective tables can be found in machine-readable json format in the supporting Information. A list of particularly important columns is given in Table S1.

Table S1: List of particularly important columns and their explanations. For a full list for all columns please see the supporting information.

| Column | Explanation |
|---|---|
| pdb | PDB id |
| name | NAOMI name, unique per PDB entry |
| skip_reason | Skip reason |
| usmiles | usmiles of the compound |
| activity.standard_value | Value of activity |
| activity.standard_relation | Relation of activity (e.g. =,>) |
| activity.standard_type | Type of activity (e.g. IC50, EC50) |
| activity.standard_units | Unit of activity (always nM for IC50,KI,KD,...) |
| assay.chembl_id | ChEMBL assay id |
| chembl_ligand.chembl_id | ChEMBL ligand id |
| target.chembl_id | ChEMBL target id |
| Target_Type | Target Type (e.g. single protein, protein familiy) |
| chembl_activity_data_validity_comment | Data validity comment in the ChEMBL |
| chembl_activity_potential_duplicate | ChEMBL potential duplicate flag |
| chembl_assay_assay_type | ChEMBL assay type (e.g. B for binding) |
| chembl_assay_confidence_score | ChEMBL assay confidence score (9 best) |
| Level_pm | Protein matching level |
| small_molecule_matching_confidence_level.comment | Molecule matching level |
| N_mutations | Number of mutations in active site |
| mw | Weight of the molecule [$\frac{g}{mol}$] |
| resolution | Resolution of the PDB structure |
| OWAB | Occupancy of weighted atomic b-factors[S1] |
| $EDIA_m$ | $EDIA_m$[S2] |
| ligand_structure | Merge indicator for ligand and structure data (left_only, right_only, both) |
| activities_structureligand | Merge indicator for activity and ligand/structure data |

#### 2 Further StrAcTable analysis

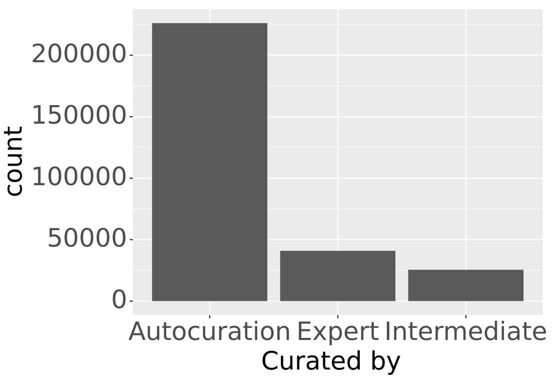


(a)

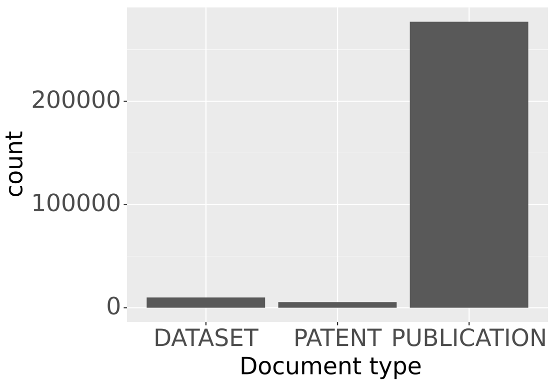


(b)

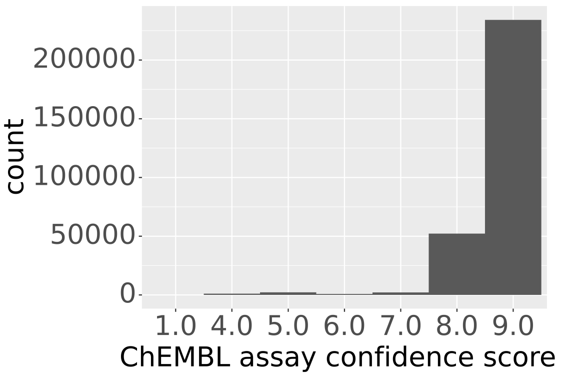


(c)

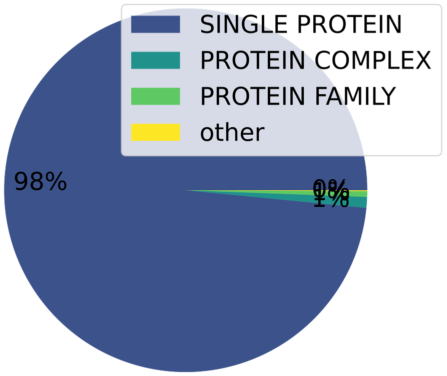


(d)

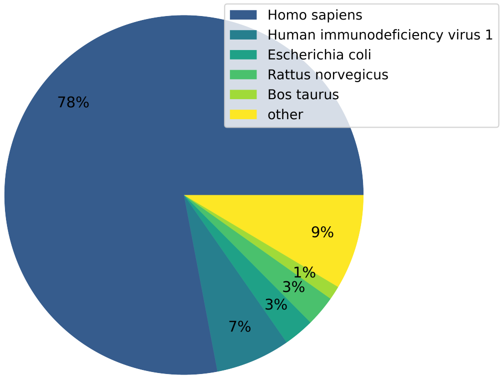


(e)

Figure S1: Analysis of the occurences of different curation strategies (a), document types (b), assay confidence scores (c), target types (d) and organisms (e) for all entries in StrAcTableF containing data from all respective tools.

### 3 LigandExtractor skip reasons

The LigandExtractor tool was developed to list all potential ligands in a PDB file and report any problems with a ligand that either arise from inconsistencies with the annotated metadata within the PDB file or from its interpretation using the NAOMI cheminformatics library.[S3,S4] It was first described in Flachsenberg et al. (2024).[S5] Table S6 of Flachsenberg et al. (2024)[S5] only contained a simple list of the occurring problems (i.e., skip reasons) along with their frequency. Here, we provide, for each possible ligand classification, at least one exemplary PDB code for further illustration. All ligands and residues are identified by the tuple `<resName>_<chainID>_<resSeq><iCode>`.

Note that the metadata within PDB files might be updated at any time in the future. Therefore, we specify for each PDB code the most recent version and revision date where the described behavior is known to occur.

**NotSkipped**

No problem was detected. An example is ligand `SOX_B_1559` in *1gm8*[S6,S7] (v1.4, 2024-05-08).

**LigandIsMetal**

The ligand is an isolated metal ion. One example is the $Ca^{2+}$ ion `CA_B_1558` in *1gm8*[S6,S7] (v1.4, 2024-05-08).

**LigandIsCovalentOrHeteroResidue**

The (potential) ligand is treated as being part of a macromolecule. This includes non-standard residues in protein chains (e.g., phosphorylated serine `SEP_A_46` in *1fu0*[S8,S9] (v1.4, 2024-10-16), and covalent ligands (e.g., ligand `FHR_A_405` in *6lze*[S10,S11] (v2.6, 2024-10-23)).

**LigandIsIncomplete**

In REMARK 470[S13] or REMARK 610,[S14] the ligand is annotated to have missing heavy atoms. One example is ligand `P33_A_201` in *4xjy*[S15] (v2.1, 2024-11-06).

**LigandContainsMetal**

The ligand molecule contains a metal atom. A common example is the iron-containing prosthetic group heme, e.g., ligand `HEM_A_143` in *1g9v*[S16,S17] (v1.4, 2023-08-09).

**LigandIsPolymerWithMissingCoordinates**

In general, oligopeptides and oligonucleotides with up to nine residues are considered to be ligand molecules. However, we check standard amino acids and nucleic acids for completeness, i.e., if any expected atom is missing. Here, either a side chain atom can be missing (e.g., ligand `ARG_P_194_ARG_P_195_ARG_P_196_GLN_P_197_TPO_P_198` in *7ort*[S18] (v1.2, 2024-11-06) is missing side chain atoms in the N-terminal arginine) or an oxygen of the C-terminal carboxy group (e.g., ligand `GLY_B_1_ALA_B_2_LYS_B_3` in *1gdn*[S19,S20] (v1.4, 2024-11-06)). The latter case also includes covalent peptide ligands, e.g., ligand `VAL_A_4_GLU_A_5_PRO_A_6_ILE_A_7` in *1h9l*[S21,S22] (v1.4, 2024-10-16), where an acyl–enzyme complex is formed.

**LigandWasSkippedByAltloc**

This is connected to the handling of alternate locations: Here, we globally use the first annotated alternate location identifier (usually 'A') for all residues. However, if a residue is exclusively annotated in another alternate location, it will not be built. An example is ligand `QTO_A_201` in *5sqm*[S23,S24] (v1.2, 2023-09-20), where the residue `QTO_A_201` is exclusively annotated with alternate location 'B'.

**LigandIsUnknownMetal**

We found a metal ion that is not given in the HET section.[S25] This can happen if there are metal-containing non-standard amino acid residues (e.g., ligand `BFD_A_170` (aspartate beryllium trifluoride) in *4xq0*[S26,S27] (v1.6, 2024-11-13) in the protein. Here, the metal is treated as a separate entity (`BE_A_170`) that is not explicitly listed in the HET section.

A similar effect can also happen for heme prosthetic groups that are covalently bound to the protein (e.g., metal `FE_A_201` in *4xxl*[S28,S29] (v2.1, 2024-10-16) that is actually part of the ligand `HEC_A_201`).

**LigandIsIncompleteNaomi**

The ligand molecule could not be correctly built by our NAOMI library,[S3,S4] and the molecule is decomposed into multiple parts. One example is ligand `BFD_A_170` in *4xq0*[S26,S27] (v1.6, 2024-11-13) where the `BFD` (aspartate beryllium trifluoride) is decomposed into the aspartate residue (part of the protein), the beryllium ion (see also 'LigandIsUnknownMetal') as well as three fluoride ions. Of the latter, only one is part of the complex.

**LigandCouldNotBeBuilt**

These molecules cannot be built by our NAOMI library[S3,S4] at all. This includes molecules with unsupported valences, e.g., bonding in carborane `34B_A_1188` in *2c2s*[S30,S31] (v1.5, 2024-05-01).

Furthermore, it also includes metal ions for which the charge could not be determined from the ATOM[S32] or HETATM[S33] section alone. One example is `TL_A_114` in *2hbn*[S34,S35] (v1.4, 2023-08-30) which could generally be $Tl^{+}$ and $Tl^{3+}$. (This ambiguity could be resolved by considering the HET code that designates the ion as $Tl^{+}$.)

**LigandIsUnknownAtom**

The ligand could not be built by the NAOMI library[S3,S4] and has the HET code `UNX`, `UNL`, or `UNK`. Here, the interpretation usually fails because the ligand only consists of an unknown atom (e.g., ligand `UNX_A_401` in *4xx6*[S36] (v2.2, 2024-10-23)).

**LigandHasUnknownHetcode**

The ligand is not a standard amino acid/nucleic acid, and it is not listed in the FORMUL section,[S37] or the given formula for the ligand is empty. An example is ligand `XWD_R_401` in *8x3l*[S38,S39] (v1.2, 2025-06-25) where the formula is empty.

**LigandIsPolymerWithChainBreak**

In general, oligopeptides and oligonucleotides with up to nine residues are considered to be ligand molecules. However, we require the oligomeric ligand to have a different chain name than the macromolecule or other oligomeric ligands (if it has the same type, i.e., nucleic acid/amino acid). This is done to ensure that there are no missing bonds, no missing residues, and that a fragment of a macromolecule is not incidentally interpreted as a ligand. An example of a chain break due to a missing bond are the two (pseudo)ligands `CYS_D_1_ASP_D_2` and `ALA_D_4_ASN_D_5_PHE_D_6_LYS_D_7` in *1p13*[S40,S41] (v1.3, 2024-12-25). These should actually form a heptapeptide with an *O*-Phosphotyrosine in between (`PTR_D_3`). However, the distances between `PTR_D_3` and `ASP_D_2`, as well as `ALA_D_4`, are both too large to actually be recognized as peptide bonds.

Similarly, the two (pseudo)ligands `LEU_J_77_SER_J_78` and `GLY_J_84_VAL_J_85` in *6gjh*[S42,S43] (v1.3, 2024-10-16) should actually form a longer peptide. Here, not only bonds, but also residues in between are missing. The latter is confirmed in the corresponding primary publication.[S43]

**LigandIsCovalentOrIncompletePolymer**

There is a connection given in the LINK section,[S44] but the residues are not attached to each other. This can, for example, be a non-detected covalent ligand, e.g., `8WY_B_1102` in *5v4q*[S45,S46] (v1.5, 2023-10-04) that should be connected to `THR_B_381` via two bonds, but the connection was not correctly interpreted by the NAOMI library.[S3,S4]

An alternative reason is an annotated LINK between a ligand and a metal ion that is never modelled by the NAOMI library,[S3,S4] e.g., ligand `COM_A_1205` in *7b2c*[S47,S48] (v1.1, 2024-01-31) is annotated to be bound to the nickel within cofactor `USN_A_1201`.

**LigandIsPolymerWithDifferentChains**

In general, oligopeptides and oligonucleotides with up to nine residues are considered to be ligand molecules. However, we check if all monomers come from the same chain. If this is not the case, we do not consider this to be a ligand. The rationale behind this filter is that usually oligomers should be defined as their own chain (see also 'LigandIsPolymerWithChainBreak').

On the one hand, this prevents molecules from being connected that obviously should not be connected: Analyzing the connection of ligands `U10_L_303` and `LDA_M_404` in *8vtj*[S49,S50] (v1.1, 2025-09-24) show unreasonable bond angles of 148 and 135 degrees at $sp^3$-carbons.

On the other hand, the assumption of homogeneous chain IDs for oligomeric ligands is not always true, especially when oligosaccharides are involved: Here, often one/the sugar part has its own chain ID, e.g., xylotriose `XYP_D_1_XYP_D_2_XYP_A_1001` in *4xuq*[S51,S52] (v2.1, 2024-01-10) or the glycosylated serine `NGA_E_1_GAL_E_2_SER_A_147` in *1y2x*[S53,S54] (v2.2, 2024-04-03).

**LigandHasUnexpectedFormula**

For a monomeric ligand, the molecular formula excluding hydrogens calculated from the molecule read by NAOMI[S3,S4] differs from the one specified in the FORMUL section.[S55] One example is ligand `4US_A_701` in *5bq0*[S56,S57] (v1.1, 2024-03-06), where in the considered alternate location 'A', the nitrogen atom 'N4' is missing, leading to a different molecular formula.

**InternalError**

Other, non-expected problems were detected. The only known case is ligand `TPO_A_158X` (and `TPO_B_158X`) in *8haw*[S58,S59] (v1.1, 2024-10-23). The problem here is an inconsistent insertion code, i.e., in the HET section,[S60] the residue is announced as `TPO_A_158X` (with upper case insertion code), but in the coordinate section, it is labeled as `TPO_A_158x` (with lower case insertion code).

**LigandIsMissingInModel1**

Table S6 in Flachsenberg et al. (2024)[S5] counted exactly one case where the PDB file contained multiple models, but the ligand in question was not part of the first model. This referred to ligand `U5P_B_903` in *2icy*.[S61,S62] However, the metadata has been corrected for the current entry version 1.4 (2023-08-30). Therefore, currently, no example for this ligand classification exists anymore.

### 4 Usage of StrAcTable

As the StrAcTable is already being used in our lab to develop further benchmarks and analysis, we want to highight some common pitfalls or usage advice to our users.

#### 4.1 Unique Keys in the StrAcTable

When working with the StrAcTable, there are many approaches which require a unique key for a protein-ligand pair. For all PDB structures, the combination of PDB-ID and Naomi ligand name (which is a combination of the ligand HETcode, chain and residue ID) describes a unique protein-ligand pair. When also taking activities into consideration, the combination of the aforementioned key with the activity id should be unique. The only exception to this rule are molecules which cannot be read correctly by Naomi like inorganic clusters. If inorganic clusters are found in the PDB (one example is the ligand JGH in the PDB ID 6qsh) they are read into multiple different molecule datastructures using NAOMI and each part of the complete molecule is handled separately for calculations like EDIAm. Thankfully, these cases as well as many other in which NAOMI either struggles to correctly model the experimental reality or the experimental model is faulty can be filtered out using the Skipreasons computed using the LigandExtractor.

#### 4.2 Handling measurements of racemic mixtures

While ligands modelled in crystal structures have to be modelled as one specific isomer, it is quite frequent to use racemic mixtures when measuring binding affinities (see Figure S2). To avoid using datapoints that have been measured using racemic mixtures, users should only use the two best ligand matching criteria (Full inchi match/ chiral usmiles match), as in these cases the identical isomer is mandatory.

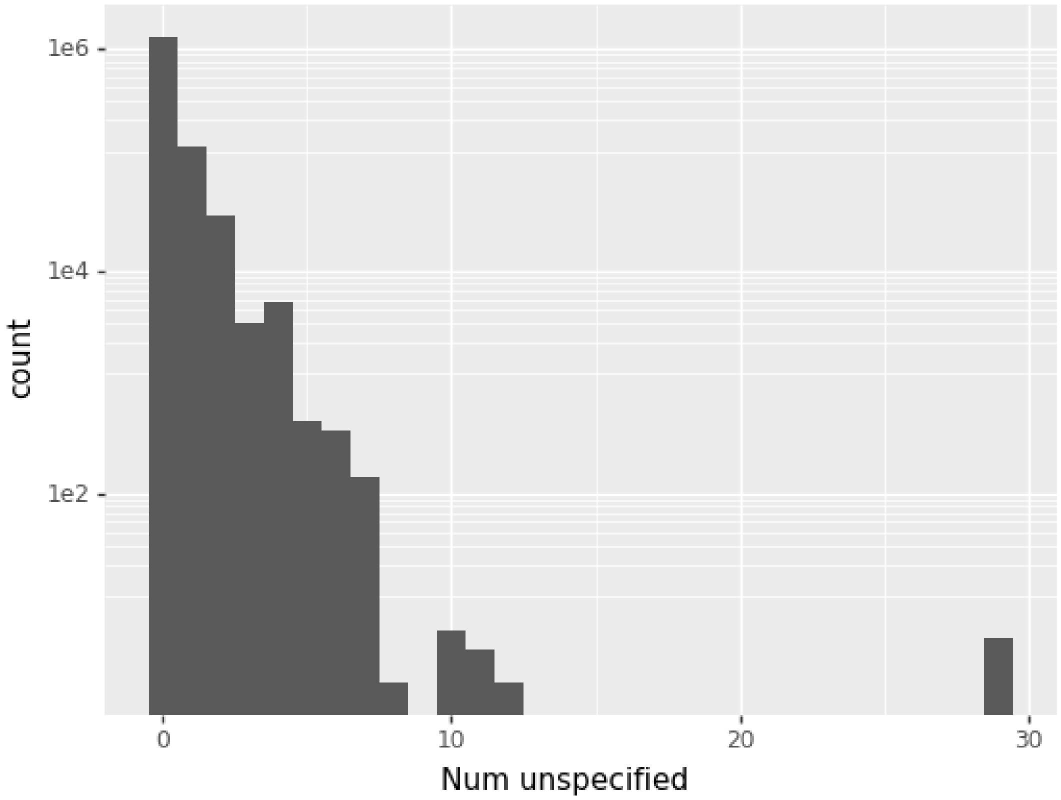


Figure S2: Number of unspecified stereocenters according to RDKit[S63] for all entries in the filtered StrAcTable that are not skipped and have activity data.

### 5 Changes in supplied electron densities

In August 2022, we learned that the electron densities supplied through EMBL-EBI had changed. In some structures, the electron density now does not encompass the complete modeled structure (e.g., PDB-ID `7AQI`[S64,S65]), or the electron density is moved in three-dimensional space compared to the modeled structure (e.g., PDB-ID `5RT2`[S66,S67]). While this can be fixed by extending and moving the electron density, this step is not initially done in the StructureProfiler. To circumvent this problem, we used GEMMI[S68] to extend and use the extended electron densities. However, for some structures (e.g., PDB-ID `1MZC`[S69,S70] or `4OCO`[S71,S72]), this results in electron densities that are not readable using the StructureProfiler due to problems of GEMMI[S68] with the dsn6 electron density format. To achieve a solution that is generalizable for all structures in the PDB, we generate the electron density from the mtz file encompassing the complete structural model using GEMMI,[S68] circumventing any issues with specific electron density file formats. However, in 2024 the mtz files were discontinued, which necessitated another change to generate the electron densities from the 2fo-fc cif files. Both approaches have been validated against the old $EDIA_m$ values before 2022. This workflow has been added to ProteinsPlus.[S73–S75]

### 6 Protein-ligand matching

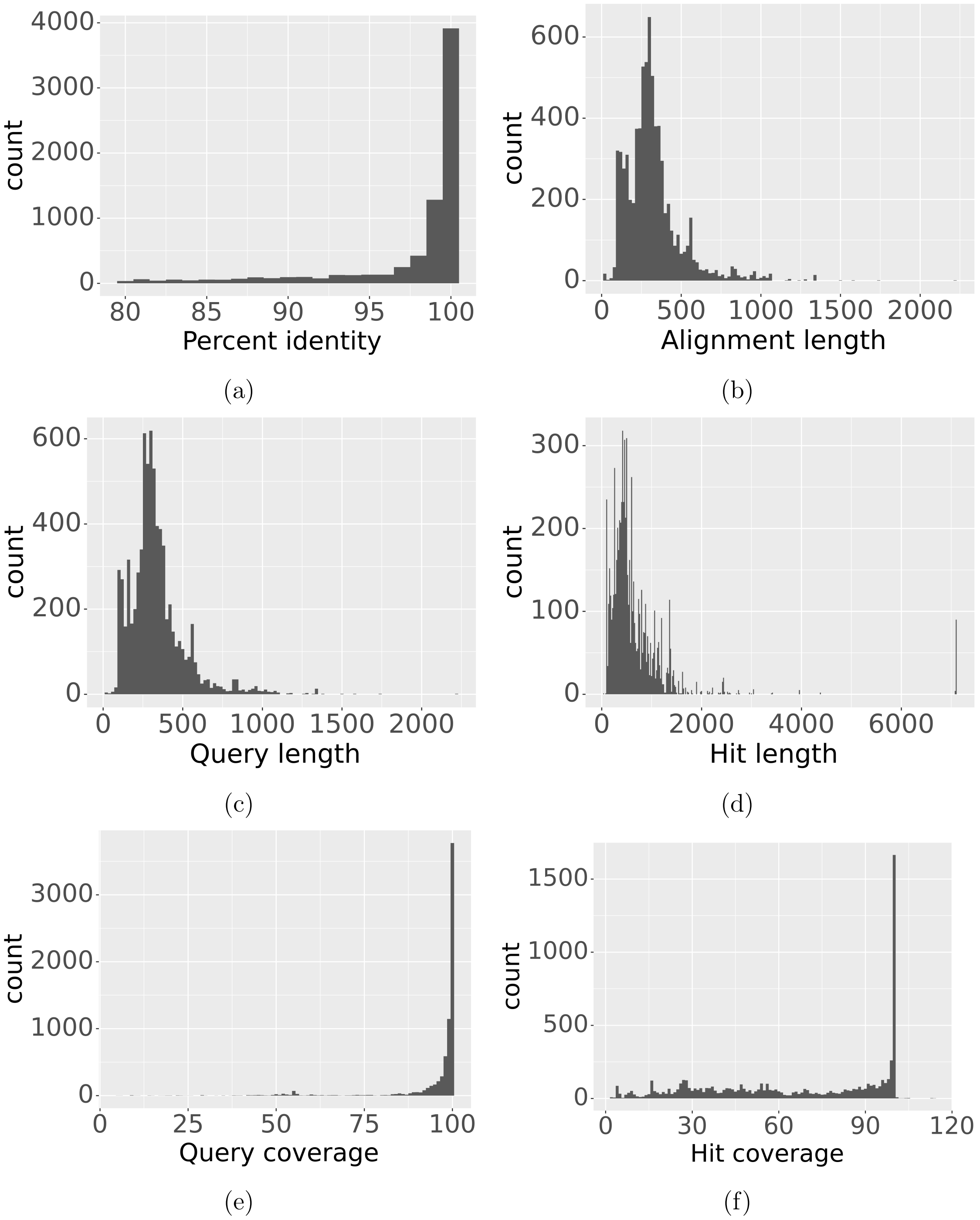


Figure S3: Histograms of the percent identity (a), alignment length (b), query length (c), hit length (d), query coverage (e), and hit coverage (f) for all unique blast match ids in the raw StrAcTable with at least one matching entry with structure and ligand data.

#### 6.1 Validation of the protein matching heuristic

ActivityFinder provides users with many possible matches to ChEMBL targets and activities for a single protein-ligand pair or PDB structure, which poses the challenge of finding the best-matching ChEMBL target for each in an automated manner. Examples of multiple ChEMBL targets for a given entry include targets from different organisms, slightly different versions of the same protein, or other target types such as single proteins, protein complexes, or protein families. The PDB structures and targets of the DUD-E[S76] dataset were used to develop and test a heuristic algorithm for automatic target selection. To compare the ability of a knowledgeable user and ActivityFinder with our heuristic to retrieve the most relevant ChEMBL target, a manual matching was conducted using metadata in the PDB and ChEMBL for each of the 102 targets and their respective structures (see Table S2). In the activity mode, no matching activities were found in 23 cases; in the target mode, two entries (Target AOFB and CASP3) produced no output, resulting in 100 possible targets in the target mode and 79 in the activity mode. In addition, for the activity mode, there are three PDBs with multiple ligands where activities can be found, meaning there are 82 total entries. Focusing on the activity mode, 73 of 82 entries (89.02%) can be correctly linked. The nine missed entries were due to activity data being measured against a slightly different target, for example, the protein of a different organism (e.g., DAY/`3BQD`[S77,S78]) or a protein complex rather than a single protein (e.g., GRIA2, `3KGC`[S79]). In each case, the manual ChEMBL target was not linked by ActivityFinder, so the heuristic could not select it. In target mode, the manually assigned target is reproduced in 96 of 100 potential cases (96.00 %). In one case, the heuristic selects a specific subunit of the protein rather than the entire complex (FNTA, `3E37`[S80,S81]). Twice (AKT1 / `3CQW`[S82,S83] and HIVPR / `1XL2`[S84,S85]), the mismatch is a failure in our heuristic, as the manual ChEMBL target and a very similar target can only be differentiated by the tiebreaker. In the other case (HIVINT / `3NF7`[S86,S87]) , the found ChEMBL target has a slightly better sequence match. Since StrAcTable will be provided both unfiltered and filtered to the best-matching ChEMBL target, users can use our validated matching or create their own algorithm.

Table S2: Detailed results of the protein matching experiment between manually matched ChEMBL targets based on metadata and the results of the ActivityFinder. Target Column describes the manually matched target, Found target describes the ActivityFinder target if they are different. Reason describes the reason for which a different target was found and Difference describes the difference between the two targets. Mode describes the mode of the ActivityFinder, either Activity mode only finding targets with ligands with matching activities or target mode ignoring data of present ligands.

| Name | PDB | Target | Found target | Reason | Difference | Type |
|---|---|---|---|---|---|---|
| AA2AR | 3eml | CHEMBL251 | | | | Activities |
| AA2AR | 3eml | CHEMBL251 | | | | Targets |
| ABL1 | 2hzi | CHEMBL1862 | | No Activities found | | Activities |
| ABL1 | 2hzi | CHEMBL1862 | | | | Targets |
| ACE | 3bkl | CHEMBL1808 | | No Activities found | | Activities |
| ACE | 3bkl | CHEMBL1808 | | | | Targets |
| ACES | 1e66 | CHEMBL4780 | | | | Activities |
| ACES | 1e66 | CHEMBL4780 | | | | Targets |
| ADA | 2e1w | CHEMBL2966 | CHEMBL1910 | Molecule not measured against perfect target | Organism difference | Activities |
| ADA | 2e1w | CHEMBL2966 | | | | Targets |
| ADA17 | 2oi0 | CHEMBL3706 | | | | Activities |
| ADA17 | 2oi0 | CHEMBL3706 | | | | Targets |
| ADRB1 | 2vt4 | CHEMBL213 | | No Activities found | | Activities |
| ADRB1 | 2vt4 | CHEMBL213 | | | | Targets |
| ADRB2 | 3ny8 | CHEMBL210 | | | | Activities |
| ADRB2 | 3ny8 | CHEMBL210 | | | | Targets |
| AKT1 | 3cqw | CHEMBL4282 | | | | Activities |
| AKT1 | 3cqw | CHEMBL4282 | CHEMBL262 | Target number is smaller | Different target | Targets |
| AKT2 | 3d0e | CHEMBL2431 | | | | Activities |
| AKT2 | 3d0e | CHEMBL2431 | | | | Targets |
| ALDR | 2hv5 | CHEMBL1900 | | | | Activities |
| ALDR | 2hv5 | CHEMBL1900 | | | | Targets |
| AMPC | 1l2s | CHEMBL2026 | | | | Activities |
| AMPC | 1l2s | CHEMBL2026 | | | | Targets |
| ANDR | 2am9 | CHEMBL1871 | | No Activities found | | Activities |
| ANDR | 2am9 | CHEMBL1871 | | | | Targets |
| AOFB | 1s3b | CHEMBL2039 | | No Activities found | | Activities |
| AOFB | 1s3b | CHEMBL2039 | | No valid ligands found | | Targets |
| BACE1 | 3l5d | CHEMBL4822 | | | | Activities |
| BACE1 | 3l5d | CHEMBL4822 | | | | Targets |
| BRAF | 3d4q | CHEMBL5145 | | | | Activities |
| BRAF | 3d4q | CHEMBL5145 | | | | Targets |

| | | | | |
|---|---|---|---|---|
| CAH2 | 1bcd | CHEMBL205 | | Activities |
| CAH2 | 1bcd | CHEMBL205 | | Targets |
| CASP3 | 2cnk | Nothing found | No Activities found | Activities |
| CASP3 | 2cnk | Nothing found | | Targets |
| CDK2 | 1h00 | CHEMBL301 | | Activities |
| CDK2 | 1h00 | CHEMBL301 | | Targets |
| COMT | 3bwm | CHEMBL2023 | No activities found | Activities |
| COMT | 3bwm | CHEMBL2023 | | Targets |
| CP2C9 | 1r9o | CHEMBL3397 | | Activities |
| CP2C9 | 1r9o | CHEMBL3397 | | Targets |
| CP3A4 | 3nxu | CHEMBL340 | | Activities |
| CP3A4 | 3nxu | CHEMBL340 | | Targets |
| CSF1R | 3krj | CHEMBL1844 | | Activities |
| CSF1R | 3krj | CHEMBL1844 | | Targets |
| CXCR4 | 3odu | CHEMBL2107 | | Activities |
| CXCR4 | 3odu | CHEMBL2107 | | Targets |
| DEF | 1lru | CHEMBL1795101 | | Activities |
| DEF | 1lru | CHEMBL1795101 | | Targets |
| DHI1 | 3frj | CHEMBL4235 | | Activities |
| DHI1 | 3frj | CHEMBL4235 | | Targets |
| DPP4 | 2i78 | CHEMBL284 | No Activities found | Activities |
| DPP4 | 2i78 | CHEMBL284 | | Targets |
| DRD3 | 3pbl | CHEMBL234 | | Activities |
| DRD3 | 3pbl | CHEMBL234 | | Targets |
| DYR | 3nxo | CHEMBL202 | | Activities |
| DYR | 3nxo | CHEMBL202 | | Targets |
| EGFR | 2rgp | CHEMBL203 | | Activities |
| EGFR | 2rgp | CHEMBL203 | | Targets |
| ESR1 | 1sj0 | CHEMBL206 | | Activities |
| ESR1 | 1sj0 | CHEMBL206 | | Targets |
| ESR2 | 2fsz | CHEMBL242 | | Activities |
| ESR2 | 2fsz | CHEMBL242 | | Targets |
| FA10 | 3kl6 | CHEMBL244 | | Activities |
| FA10 | 3kl6 | CHEMBL244 | | Targets |
| FA7 | 1w7x | CHEMBL3991 | | Activities |
| FA7 | 1w7x | CHEMBL3991 | | Targets |
| FABP4 | 2nnq | CHEMBL2083 | | Activities |
| FABP4 | 2nnq | CHEMBL2083 | | Targets |
| FAK1 | 3bz3 | CHEMBL2695 | | Activities |
| FAK1 | 3bz3 | CHEMBL2695 | | Targets |

| | | | | | | |
|---|---|---|---|---|---|---|
| FGFR1 | 3c4f | CHEMBL3650 | | | | Activities |
| FGFR1 | 3c4f | CHEMBL3650 | | | | Targets |
| FKB1A | 1j4h | CHEMBL1902 | | No Activities found | | Activities |
| FKB1A | 1j4h | CHEMBL1902 | | | | Targets |
| FNTA | 3e37 | CHEMBL2094108 | | No Activities found | | Activities |
| FNTA | 3e37 | CHEMBL2094108 | CHEMBL271 | Target type | Subunit of whole protein | Targets |
| FPPS | 1zw5 | CHEMBL1782 | | | | Activities |
| FPPS | 1zw5 | CHEMBL1782 | | | | Targets |
| GCR | 3bqd | CHEMBL2034 | CHEMBL3368 | Molecule not measured against perfect target | Organism difference | Activities |
| GCR | 3bqd | CHEMBL2034 | | | | Targets |
| GLCM | 2v3f | CHEMBL2179 | | No Activities found | | Activities |
| GLCM | 2v3f | CHEMBL2179 | | | | Targets |
| GRIA2 | 3kgc | CHEMBL3503 | CHEMBL4016/CHEMBL2093871 | Molecule not measured against perfect target | Organism difference/ Protein Complex instead of single Protein | Activities |
| GRIA2 | 3kgc | CHEMBL3503 | | | | Targets |
| GRIK1 | 1vso | CHEMBL2919 | | | | Activities |
| GRIK1 | 1vso | CHEMBL2919 | | | | Targets |
| HDAC2 | 3max | CHEMBL1937 | | | | Activities |
| HDAC2 | 3max | CHEMBL1937 | | | | Targets |
| HDAC8 | 3f07 | CHEMBL3192 | CHEMBL2093865 | Molecules not measured against perfect target | Protein Familiy instead of single Protein | Activities |
| HDAC8 | 3f07 | CHEMBL3192 | | | | Targets |
| HIVINT | 3nf7 | CHEMBL3471 | | No Activities found | | Activities |
| HIVINT | 3nf7 | CHEMBL3471 | CHEMBL2366505 | Sequence matching | Almost identical target | Targets |
| HIVPR | 1xl2 | CHEMBL3638326 | CHEMBL243 | | Almost identical target | Activities |
| HIVPR | 1xl2 | CHEMBL3638326 | CHEMBL5823 | Target number is smaller | | Targets |
| HIVRT | 3lan | CHEMBL247 | | No Activities found | | Activities |
| HIVRT | 3lan | CHEMBL247 | | | | Targets |
| HMDH | 3ccw | CHEMBL402 | | | | Activities |
| HMDH | 3ccw | CHEMBL402 | | | | Targets |
| HS90A | 1uyg | CHEMBL3880 | CHEMBL2095165 | Molecules not measured against perfect target | Protein Familiy instead of single Protein | Activities |
| HS90A | 1uyg | CHEMBL3880 | | | | Targets |
| HXK4 | 3f9m | CHEMBL3820 | | | | Activities |
| HXK4 | 3f9m | CHEMBL3820 | | | | Targets |
| IGF1R | 2oj9 | CHEMBL1957 | | | | Activities |
| IGF1R | 2oj9 | CHEMBL1957 | | | | Targets |
| INHA | 4trj | CHEMBL1849 | | | | Activities |
| INHA | 4trj | CHEMBL1849 | | | | Targets |
| ITAL | 2ica | CHEMBL1803 | | | | Activities |
| ITAL | 2ica | CHEMBL1803 | | | | Targets |
| JAK2 | 3lpb | CHEMBL2971 | | | | Activities |
| JAK2 | 3lpb | CHEMBL2971 | | | | Targets |

| | | | | | | |
|---|---|---|---|---|---|---|
| KIF11 | 3cjo | CHEMBL4581 | | | | Activities |
| KIF11 | 3cjo | CHEMBL4581 | | | | Targets |
| KIT | 3g0e | CHEMBL1936 | | No Activities found | | Activities |
| KIT | 3g0e | CHEMBL1936 | | | | Targets |
| KITH | 2b8t | CHEMBL1075130 | | | | Activities |
| KITH | 2b8t | CHEMBL1075130 | | | | Targets |
| KPCB | 2i0e | CHEMBL3045 | | No Activities found | | Activities |
| KPCB | 2i0e | CHEMBL3045 | | | | Targets |
| LCK | 2of2 | CHEMBL258 | | | | Activities |
| LCK | 2of2 | CHEMBL258 | | | | Targets |
| LKHA4 | 3chp | CHEMBL4618 | | | | Activities |
| LKHA4 | 3chp | CHEMBL4618 | | | | Targets |
| MAPK2 | 3m2w | CHEMBL2208 | | | | Activities |
| MAPK2 | 3m2w | CHEMBL2208 | | | | Targets |
| MCR | 2aa2 | CHEMBL1994 | | No Activities found | | Activities |
| MCR | 2aa2 | CHEMBL1994 | | | | Targets |
| MET | 3lq8 | CHEMBL3717 | | | | Activities |
| MET | 3lq8 | CHEMBL3717 | | | | Targets |
| MK01 | 2ojg | CHEMBL4040 | | | | Activities |
| MK01 | 2ojg | CHEMBL4040 | | | | Targets |
| MK10 | 2zdt | CHEMBL2637 | CHEMBL2276 | Molecules not measured against perfect target | kinase 1 instead 3 | Activities |
| MK10 | 2zdt | CHEMBL2637 | | | | Targets |
| MK14 | 2qd9 | CHEMBL260 | | | | Activities |
| MK14 | 2qd9 | CHEMBL260 | | | | Targets |
| MMP13 | 830c | CHEMBL280 | | | | Activities |
| MMP13 | 830c | CHEMBL280 | | | | Targets |
| MP2K1 | 3eqh | CHEMBL3587 | | | | Activities |
| MP2K1 | 3eqh | CHEMBL3587 | | | | Targets |
| NOS1 | 1qw6 | CHEMBL3048 | | | | Activities |
| NOS1 | 1qw6 | CHEMBL3048 | | | | Targets |
| NRAM | 1b9v | CHEMBL3377 | | | | Activities |
| NRAM | 1b9v | CHEMBL3377 | | | | Targets |
| PA2GA | 1kvo | CHEMBL3474 | | | | Activities |
| PA2GA | 1kvo | CHEMBL3474 | | | | Targets |
| PARP1 | 3l3m | CHEMBL3105 | | | | Activities |
| PARP1 | 3l3m | CHEMBL3105 | | | | Targets |
| PDE5A | 1udt | CHEMBL1827 | | | | Activities |
| PDE5A | 1udt | CHEMBL1827 | | | | Targets |
| PGH1 | 2oyu | CHEMBL2949 | | | | Activities |
| PGH1 | 2oyu | CHEMBL2949 | | | | Targets |

| | | | | |
|---|---|---|---|---|
| PGH2 | 3ln1 | CHEMBL4321 | | Activities |
| PGH2 | 3ln1 | CHEMBL4321 | | Targets |
| PLK1 | 2owb | CHEMBL3024 | | Activities |
| PLK1 | 2owb | CHEMBL3024 | | Targets |
| PNPH | 3bgs | CHEMBL4338 | | Activities |
| PNPH | 3bgs | CHEMBL4338 | | Targets |
| PPARA | 2p54 | CHEMBL239 | | Activities |
| PPARA | 2p54 | CHEMBL239 | | Targets |
| PPARD | 2znp | CHEMBL3979 | | Activities |
| PPARD | 2znp | CHEMBL3979 | | Targets |
| PPARG | 2gtk | CHEMBL235 | | Activities |
| PPARG | 2gtk | CHEMBL235 | | Targets |
| PRGR | 3kba | CHEMBL208 | | Activities |
| PRGR | 3kba | CHEMBL208 | | Targets |
| PTN1 | 2azr | CHEMBL335 | No Activities found | Activities |
| PTN1 | 2azr | CHEMBL335 | | Targets |
| PUR2 | 1njs | CHEMBL3972 | No Activities found | Activities |
| PUR2 | 1njs | CHEMBL3972 | | Targets |
| PYGM | 1c8k | CHEMBL4696 | | Activities |
| PYGM | 1c8k | CHEMBL4696 | | Targets |
| PYRD | 1d3g | CHEMBL1966 | | Activities |
| PYRD | 1d3g | CHEMBL1966 | | Targets |
| RENI | 3g6z | CHEMBL286 | | Activities |
| RENI | 3g6z | CHEMBL286 | | Targets |
| ROCK1 | 2etr | CHEMBL3231 | | Activities |
| ROCK1 | 2etr | CHEMBL3231 | | Targets |
| RXRA | 1mv9 | CHEMBL2061 | | Activities |
| RXRA | 1mv9 | CHEMBL2061 | | Targets |
| SAHH | 1li4 | CHEMBL2664 | | Activities |
| SAHH | 1li4 | CHEMBL2664 | | Targets |
| SRC | 3el8 | CHEMBL3655 | No Activities found | Activities |
| SRC | 3el8 | CHEMBL3655 | | Targets |
| TGFR1 | 3hmm | CHEMBL4439 | | Activities |
| TGFR1 | 3hmm | CHEMBL4439 | | Targets |
| THB | 1q4x | CHEMBL1947 | No Activities found | Activities |
| THB | 1q4x | CHEMBL1947 | | Targets |
| THRB | 1ype | CHEMBL204 | No Activities found | Activities |
| THRB | 1ype | CHEMBL204 | | Targets |
| TRY1 | 2ayw | CHEMBL3769 | No Activities found | Activities |
| TRY1 | 2ayw | CHEMBL3769 | | Targets |

| | | | | | | |
|---|---|---|---|---|---|---|
| TRYB1 | 2zec | CHEMBL2617 | | No Activities found | | Activities |
| TRYB1 | 2zec | CHEMBL2617 | | | | Targets |
| TYSY | 1syn | CHEMBL2555 | CHEMBL6137 | Molecule not measured against perfect target | Organism difference | Activities |
| TYSY | 1syn | CHEMBL2555 | | | | Targets |
| UROK | 1sqt | CHEMBL3286 | | | | Activities |
| UROK | 1sqt | CHEMBL3286 | | | | Targets |
| VGFR2 | 2p2i | CHEMBL279 | | | | Activities |
| VGFR2 | 2p2i | CHEMBL279 | | | | Targets |
| WEE1 | 3biz | CHEMBL5491 | | | | Activities |
| WEE1 | 3biz | CHEMBL5491 | | | | Targets |
| XIAP | 3hl5 | CHEMBL4198 | | | | Activities |
| XIAP | 3hl5 | CHEMBL4198 | | | | Targets |

(S8) Audette, G. F.; Engelmann, R.; Hengstenberg, W.; Deutscher, J.; Hayakawa, K.; Quail, J. W.; Delbaere, L. T. J. CRYSTAL STRUCTURE ANALYSIS OF THE

PHOSPHO-SERINE 46 HPR FROM ENTEROCOCCUS FAECALIS. 2000; https://doi.org/10.2210/pdb1FU0/pdb.

(S9) Audette, G. F.; Engelmann, R.; Hengstenberg, W.; Deutscher, J.; Hayakawa, K.; Quail, J. W.; Delbaere, L. T. J. The 1.9 Å Resolution Structure of Phospho-serine 46 HPr from *Enterococcus faecalis*. *Journal of Molecular Biology* **2000**, *303*, 545–553.

(S10) Zhang, B.; Zhang, Y.; Jing, Z.; Liu, X.; Yang, H.; Liu, H.; Rao, Z.; Jiang, H. The crystal structure of COVID-19 main protease in complex with an inhibitor 11a. 2020; https://doi.org/10.2210/pdb6LZE/pdb.

(S11) Dai, W.; Zhang, B.; Jiang, X.-M.; Su, H.; Li, J.; Zhao, Y.; Xie, X.; Jin, Z.; Peng, J.; Liu, F.; Li, C.; Li, Y.; Bai, F.; Wang, H.; Cheng, X.; Cen, X.; Hu, S.; Yang, X.; Wang, J.; Liu, X.; Xiao, G.; Jiang, H.; Rao, Z.; Zhang, L.-K.; Xu, Y.; Yang, H.; Liu, H. Structure-based design of antiviral drug candidates targeting the SARS-CoV-2 main protease. *Science* **2020**, *368*, 1331–1335.

(S12) Protein Data Bank Contents Guide: Atomic Coordinate Entry Format Description Version 3.30. https://files.wwpdb.org/pub/pdb/doc/format_descriptions/Format_v33_Letter.pdf, accessed on 10/27/2025.

(S13) Ref. S12, pp. 127–128.

(S14) Ref. S12, p. 138.

(S15) Kauer, S.; Schirmer, T. Periplasmic repressor protein YfiR. 2015; https://doi.org/10.2210/pdb4XJY/pdb.

(S16) Safo, M. K.; Moure, C. M.; Burnett, J. C.; Joshi, G. S.; Abraham, D. J. HIGH RESOLUTION CRYSTAL STRUCTURE OF DEOXY HEMOGLOBIN COMPLEXED WITH A POTENT ALLOSTERIC EFFECTOR. 2000; https://doi.org/10.2210/pdb1G9V/pdb.

(S17) Safo, M. K.; Moure, C. M.; Burnett, J. C.; Joshi, G. S.; Abraham, D. J. High-resolution crystal structure of deoxy hemoglobin complexed with a potent allosteric effector. *Protein Science* **2001**, *10*, 951–957.

(S18) Centorrino, F.; Wu, Q.; Ottmann, C. Ternary complex of 14-3-3 sigma, p27pT198 phosphopeptide, and WQ176. 2021; https://doi.org/10.2210/pdb7ORT/pdb.

(S19) Rypniewsky, W. R.; Oestergaard, P.; Noerregaard-Madsen, M.; Dauter, M.; Wilson, K. S. FUSARIUM OXYSPORUM TRYPSIN AT ATOMIC RESOLUTION. 2000; https://doi.org/10.2210/pdb1GDN/pdb.

(S20) Rypniewski, W. R.; Østergaard, P. R.; Nørregaard-Madsen, M.; Dauter, M.; Wilson, K. S. *Fusarium oxysporum* trypsin at atomic resolution at 100 and 283 K: a study of ligand binding. *Acta Crystallographica Section D: Biological Crystallography* **2001**, *57*, 8–19.

(S21) Wright, P. A.; Wilmouth, R. C.; Clifton, I. J.; Schofield, C. J. PORCINE PANCREATIC ELASTASE COMPLEXED WITH ACETYL-VAL-GLU-PRO-ILE-COOH. 2001; https://doi.org/10.2210/pdb1H9L/pdb.

(S22) Wright, P. A.; Wilmouth, R. C.; Clifton, I. J.; Schofield, C. J. Kinetic and crystallographic analysis of complexes formed between elastase and peptides from β-casein. *European Journal of Biochemistry* **2001**, *268*, 2969–2974.

(S23) Correy, G. J.; Fraser, J. S. PanDDA analysis group deposition -- Crystal structure of SARS-CoV-2 NSP3 macrodomain in complex with Z5028367849 - (S) isomer. 2022; https://doi.org/10.2210/pdb5SQM/pdb.

(S24) Gahbauer, S.; Correy, G. J.; Schuller, M.; Ferla, M. P.; Doruk, Y. U.; Rachman, M.; Wu, T.; Diolaiti, M.; Wang, S.; Neitz, R. J.; Fearon, D.; Radchenko, D. S.; Moroz, Y. S.; Irwin, J. J.; Renslo, A. R.; Taylor, J. C.; Gestwicki, J. E.; von Delft, F.;

Ashworth, A.; Ahel, I.; Shoichet, B. K.; Fraser, J. S. Iterative computational design and crystallographic screening identifies potent inhibitors targeting the Nsp3 macrodomain of SARS-CoV-2. *Proceedings of the National Academy of Sciences* **2023**, *120*, e2212931120.

(S25) Ref. S12, pp. 159–160.

(S26) Ghosh, A.; Lima, C. D. Structure of fission yeast RNA polymerase II CTD phosphatase Fcp1-R271A bound to beryllium fluoride. 2015; `https://doi.org/10.2210/pdb4XQ0/pdb`.

(S27) Schwer, B.; Ghosh, A.; Sanchez, A. M.; Lima, C. D.; Shuman, S. Genetic and structural analysis of the essential fission yeast RNA polymerase II CTD phosphatase Fcp1. *RNA* **2015**, *21*, 1135–1146.

(S28) Beckwith, C. R.; Edwards, M. J.; Clarke, T. Crystal structure of Class 1 cytochrome MtoD from Sideroxydans lithotrophicus ES-1. 2015; `https://doi.org/10.2210/pdb4XXL/pdb`.

(S29) Beckwith, C. R.; Edwards, M. J.; Lawes, M.; Shi, L.; Butt, J. N.; Richardson, D. J.; Clarke, T. A. Characterization of MtoD from *Sideroxydans lithotrophicus:* a cytochrome c electron shuttle used in lithoautotrophic growth. *Frontiers in Microbiology* **2015**, *6*, 332.

(S30) Leung, A. K. W.; Reynolds, R. C.; Riordan, J. M.; Borhani, D. W. Human Dihydrofolate Reductase Complexed With NADPH and 2,4-Diamino-5-(1-o-carboranylmethyl)-6-methylpyrimidine, A novel boron containing, nonclassical Antifolate. 2005; `https://doi.org/10.2210/pdb2C2S/pdb`.

(S31) Reynolds, R. C.; Campbell, S. R.; Fairchild, R. G.; Kisliuk, R. L.; Micca, P. L.; Queener, S. F.; Riordan, J. M.; Sedwick, W. D.; Waud, W. R.; Leung, A. K. W.;

Dixon, R. W.; Suling, W. J.; Borhani, D. W. Novel Boron-Containing, Nonclassical Antifolates: Synthesis and Preliminary Biological and Structural Evaluation. *Journal of Medicinal Chemistry* **2007**, *50*, 3283–3289.

(S32) Ref. S12, pp. 187–189.

(S33) Ref. S12, pp. 194–195.

(S34) Gill, M. L.; Strobel, S. A.; Loria, J. P. Crystallization of the Tl+-form of the Oxytricha nova G-quadruplex. 2006; https://doi.org/10.2210/pdb2HBN/pdb.

(S35) Gill, M. L.; Strobel, S. A.; Loria, J. P. Crystallization and characterization of the thallium form of the *Oxytricha nova* G-quadruplex. *Nucleic Acids Research* **2006**, *34*, 4506–4514.

(S36) Stogios, P. J.; Nocek, B.; Xu, X.; Cui, H.; Lowden, M.; Savchenko, A. Crystal structure of a glycosylated endo-beta-1,4-xylanase (glycoside hydrolase family 10/GH10) enzyme from Gloeophyllum trabeum. 2015; https://doi.org/10.2210/pdb4XX6/pdb.

(S37) Ref. S12, pp. 164–165.

(S38) Shen, S. Y.; Yang, Z. Q.; Shao, Z. H. Cryo-EM structure of CB2-G protein complex. 2023; https://doi.org/10.2210/pdb8X3L/pdb.

(S39) Yang, X.; Zhou, P.; Shen, S.; Hu, Q.; Tian, C.; Xia, A.; Wang, Y.; Yang, Z.; Nan, J.; Zhou, Y.; Chen, S.; Tian, X.; Wu, C.; Lin, G.; Zhang, L.; Wang, K.; Zheng, T.; Zou, J.; Yan, W.; Shao, Z.; Yang, S. Entropy drives the ligand recognition in G-protein-coupled receptor subtypes. *Proceedings of the National Academy of Sciences* **2024**, *121*, e2401091121.

(S40) Sonnenburg, E. D.; Bilwes, A.; Hunter, T.; Noel, J. P. Crystal Structure of the Src SH2 Domain Complexed with Peptide (SDpYANFK). 2003; https://doi.org/10.2210/pdb1P13/pdb.

(S41) Sonnenburg, E. D.; Bilwes, A.; Hunter, T.; Noel, J. P. The Structure of the Membrane Distal Phosphatase Domain of RPTP$\alpha$ Reveals Interdomain Flexibility and an SH2 Domain Interaction Region. *Biochemistry* **2003**, *42*, 7904–7914.

(S42) Collier, M. P.; Benesch, J. L. P. Human Hsp27 (HspB1) alpha-crystallin domain in complex with a peptide mimic of its phosphorylatable N-terminal region. 2018; `https://doi.org/10.2210/pdb6GJH/pdb`.

(S43) Collier, M. P.; Alderson, T. R.; de Villiers, C. P.; Nicholls, D.; Gastall, H. Y.; Allison, T. M.; Degiacomi, M. T.; Jiang, H.; Mlynek, G.; Fürst, D. O.; van der Ven, P. F. M.; Djinovic-Carugo, K.; Baldwin, A. J.; Watkins, H.; Gehmlich, K.; Benesch, J. L. P. HspB1 phosphorylation regulates its intramolecular dynamics and mechanosensitive molecular chaperone interaction with filamin C. *Science Advances* **2019**, *5*, eaav8421.

(S44) Ref. S12, pp. 173–174.

(S45) Terzyan, S.; Hanigan, M. Crystal Structure of human GGT1 in complex with DON. 2017; `https://doi.org/10.2210/pdb5V4Q/pdb`.

(S46) Terzyan, S. S.; Cook, P. F.; Heroux, A.; Hanigan, M. H. Structure of 6-diazo-5-oxo-norleucine-bound human gamma-glutamyl transpeptidase 1, a novel mechanism of inactivation. *Protein Science* **2017**, *26*, 1196–1205.

(S47) Wagner, T.; Lemaire, O. N.; Engilberge, S. Crystal structure of the ethyl-coenzyme M reductase from Candidatus Ethanoperedens thermophilum gassed with xenon. 2020; `https://doi.org/10.2210/pdb7B2C/pdb`.

(S48) Hahn, C. J.; Lemaire, O. N.; Kahnt, J.; Engilberge, S.; Wegener, G.; Wagner, T. Crystal structure of a key enzyme for anaerobic ethane activation. *Science* **2021**, *373*, 118–121.

(S49) Mathews, I.; Tran, K.; Boxer, S. G. Crystal structure of R. sphaeroides Photosynthetic Reaction Center variant Y(M210)2-cyanophenylalanine. 2024; https://doi.org/10.2210/pdb8VTJ/pdb.

(S50) Tran, K. N.; Faries, K. M.; Magdaong, N. C. M.; Mathews, I. I.; Weaver, J. B.; Kirsh, J. M.; Holten, D.; Kirmaier, C.; Boxer, S. G. Application of Amber Suppression To Study the Role of Tyr M210 in Electron Transfer in *Rhodobacter sphaeroides* Photosynthetic Reaction Centers. *The Journal of Physical Chemistry B* **2025**, *129*, 3317–3333.

(S51) Sainz-Polo, M. A.; Sanz-Aparicio, J. Structure of the CBM22-2 xylan-binding domain from Paenibacillus barcinonensis Xyn10C in complex with xylotriose. 2015; https://doi.org/10.2210/pdb4XUQ/pdb.

(S52) Sainz-Polo, M. A.; González, B.; Menéndez, M.; Pastor, F. I. J.; Sanz-Aparicio, J. Exploring Multimodularity in Plant Cell Wall Deconstruction: STRUCTURAL AND FUNCTIONAL ANALYSIS OF Xyn10C CONTAINING THE CBM22-1–CBM22-2 TANDEM. *Journal of Biological Chemistry* **2015**, *290*, 17116–17130.

(S53) Carrizo, M. E.; Capaldi, S.; Perduca, M.; Irazoqui, F. J.; Nores, G. A.; Monaco, H. L. Crystal structure of the tetragonal form of the common edible mushroom (Agaricus bisporus) lectin in complex with T-antigen and N-acetylglucosamine. 2004; https://doi.org/10.2210/pdb1Y2X/pdb.

(S54) Carrizo, M. E.; Capaldi, S.; Perduca, M.; Irazoqui, F. J.; Nores, G. A.; Monaco, H. L. The Antineoplastic Lectin of the Common Edible Mushroom (*Agaricus bisporus*) Has Two Binding Sites, Each Specific for a Different Configuration at a Single Epimeric Hydroxyl. *Journal of Biological Chemistry* **2005**, *280*, 10614–10623.

(S55) Ref. S12, pp. 164–165.

(S56) Muckelbauer, J. K. Crystal structure of bruton agammaglobulinemia tyrosine kinase complexed with BMS-824171 AKA 6-[(3R)-3-(4-tert-bu tylbenzamido)piperidin-1-yl]-2-{[4-(morpholine-4-carbonyl) phenyl]amino}pyridine-3-carboxamide. 2015; https://doi.org/10.2210/pdb5BQ0/pdb.

(S57) Liu, Q.; Batt, D. G.; Lippy, J. S.; Surti, N.; Tebben, A. J.; Muckelbauer, J. K.; Chen, L.; An, Y.; Chang, C.; Pokross, M.; Yang, Z.; Wang, H.; Burke, J. R.; Carter, P. H.; Tino, J. A. Design and synthesis of carbazole carboxamides as promising inhibitors of Bruton's tyrosine kinase (BTK) and Janus kinase 2 (JAK2). *Bioorganic & Medicinal Chemistry Letters* **2015**, *25*, 4265–4269.

(S58) Zhao, F.; Fan, R. Y.; Guan, Z. Y.; Guo, L.; Yin, P. An auto-activation mechanism of plant non-specific phospholipase C. 2022; https://doi.org/10.2210/pdb8HAW/pdb.

(S59) Fan, R.; Zhao, F.; Gong, Z.; Chen, Y.; Yang, B.; Zhou, C.; Zhang, J.; Du, Z.; Wang, X.; Yin, P.; Guo, L.; Liu, Z. Insights into the mechanism of phospholipid hydrolysis by plant non-specific phospholipase C. *Nature Communications* **2023**, *14*, 194.

(S60) Ref. S12, pp. 159–160.

(S61) McCoy, J. G.; Wesenberg, G. E.; Phillips Jr., G. N.; Bitto, E.; Bingman, C. A.; Center for Euraryotic Structural Genomics (CESG)., Crystal Structure of a Putative UDP-glucose Pyrophosphorylase from Arabidopsis Thaliana with Bound UDP-glucose. 2006; https://doi.org/10.2210/pdb2ICY/pdb.

(S62) McCoy, J. G.; Bitto, E.; Bingman, C. A.; Wesenberg, G. E.; Bannen, R. M.; Kondrashov, D. A.; Phillips Jr, G. N. Structure and Dynamics of UDP–Glucose Pyrophosphorylase from *Arabidopsis thaliana* with Bound UDP–Glucose and UTP. *Journal of Molecular Biology* **2007**, *366*, 830–841.

(S63) RDKit: Open-source Cheminformatics. http://www.rdkit.org.

(S64) Koua, F.; Guenther, S.; Reinke, P.; Oberthuer, D.; Yefanov, O.; Gelisio, L.; Ginn, H.; Lieske, J.; Ewert, W.; Domaracky, M.; Brehm, W.; Rahmani Mashour, A.; White, T.; Knoska, J.; Pena Esperanza, G.; Tolstikova, A.; Groessler, M.; Fischer, P.; Hennicke, V.; Fleckenstein, H.; Trost, F.; Galchenkova, M.; Gevorkov, Y.; Li, C.; Awel, S.; Paulraj, L.; Ullah, N.; Falke, S.; Alves Franca, B.; Schwinzer, M.; Brognaro, H.; Werner, N.; Perbandt, M.; Tidow, H.; Seychell, B.; Beck, T.; Meier, S.; Doyle, J.; Giseler, H.; Melo, D.; Dunkel, I.; Lane, T.; Peck, A.; Saouane, S.; Hakanpaeae, J.; Meyer, J.; Noei, H.; Gribbon, P.; Ellinger, B.; Kuzikov, M.; Wolf, M.; Zhang, L.; Ehrt, C.; Pletzer-Zelgert, J.; Wollenhaupt, J.; Feiler, C.; Weiss, M.; Schulz, E.; Mehrabi, P.; Norton-Baker, B.; Schmidt, C.; Lorenzen, K.; Schubert, R.; Han, H.; Chari, A.; Fernandez Garcia, Y.; Turk, D.; Hilgenfeld, R.; Rarey, M.; Zaliani, A.; Chapman, H.; Pearson, A.; Betzel, C.; Meents, A. Structure of SARS-CoV-2 Main Protease Bound to Ifenprodil: 7aqi. 2020; https://www.wwpdb.org/pdb?id=pdb_00007aqi.

(S65) Günther, S.; Reinke, P. Y. A.; Fernández-García, Y.; Lieske, J.; Lane, T. J.; Ginn, H. M.; Koua, F. H. M.; Ehrt, C.; Ewert, W.; Oberthuer, D.; Yefanov, O.; Meier, S.; Lorenzen, K.; Krichel, B.; Kopicki, J.-D.; Gelisio, L.; Brehm, W.; Dunkel, I.; Seychell, B.; Gieseler, H.; Norton-Baker, B.; Escudero-Pérez, B.; Domaracky, M.; Saouane, S.; Tolstikova, A.; White, T. A.; Hänle, A.; Groessler, M.; Fleckenstein, H.; Trost, F.; Galchenkova, M.; Gevorkov, Y.; Li, C.; Awel, S.; Peck, A.; Barthelmess, M.; Schlünzen, F.; Lourdu Xavier, P.; Werner, N.; Andaleeb, H.; Ullah, N.; Falke, S.; Srinivasan, V.; França, B. A.; Schwinzer, M.; Brognaro, H.; Rogers, C.; Melo, D.; Zaitseva-Doyle, J. J.; Knoska, J.; Peña-Murillo, G. E.; Mashhour, A. R.; Hennicke, V.; Fischer, P.; Hakanpää, J.; Meyer, J.; Gribbon, P.; Ellinger, B.; Kuzikov, M.; Wolf, M.; Beccari, A. R.; Bourenkov, G.; Von Stetten, D.; Pompidor, G.; Bento, I.; Panneerselvam, S.; Karpics, I.; Schneider, T. R.; Garcia-Alai, M. M.; Niebling, S.; Günther, C.; Schmidt, C.; Schubert, R.; Han, H.; Boger, J.; Monteiro, D. C. F.; Zhang, L.;

Sun, X.; Pletzer-Zelgert, J.; Wollenhaupt, J.; Feiler, C. G.; Weiss, M. S.; Schulz, E.-C.; Mehrabi, P.; Karničar, K.; Usenik, A.; Loboda, J.; Tidow, H.; Chari, A.; Hilgenfeld, R.; Uetrecht, C.; Cox, R.; Zaliani, A.; Beck, T.; Rarey, M.; Günther, S.; Turk, D.; Hinrichs, W.; Chapman, H. N.; Pearson, A. R.; Betzel, C.; Meents, A. X-Ray Screening Identifies Active Site and Allosteric Inhibitors of SARS-CoV-2 Main Protease. *Science* **2021**, *372*, 642–646.

(S66) Correy, G.; Young, I.; Thompson, M.; Fraser, J. PanDDA Analysis Group Deposition – Crystal Structure of SARS-CoV-2 NSP3 Macrodomain in Complex with ZINC000008652361: 5rt2. 2020; https://www.wwpdb.org/pdb?id=pdb_00005rt2.

(S67) Schuller, M.; Correy, G. J.; Gahbauer, S.; Fearon, D.; Wu, T.; Díaz, R. E.; Young, I. D.; Carvalho Martins, L.; Smith, D. H.; Schulze-Gahmen, U.; Owens, T. W.; Deshpande, I.; Merz, G. E.; Thwin, A. C.; Biel, J. T.; Peters, J. K.; Moritz, M.; Herrera, N.; Kratochvil, H. T.; QCRG Structural Biology Consortium,; Aimon, A.; Bennett, J. M.; Brandao Neto, J.; Cohen, A. E.; Dias, A.; Douangamath, A.; Dunnett, L.; Fedorov, O.; Ferla, M. P.; Fuchs, M. R.; Gorrie-Stone, T. J.; Holton, J. M.; Johnson, M. G.; Krojer, T.; Meigs, G.; Powell, A. J.; Rack, J. G. M.; Rangel, V. L.; Russi, S.; Skyner, R. E.; Smith, C. A.; Soares, A. S.; Wierman, J. L.; Zhu, K.; O'Brien, P.; Jura, N.; Ashworth, A.; Irwin, J. J.; Thompson, M. C.; Gestwicki, J. E.; Von Delft, F.; Shoichet, B. K.; Fraser, J. S.; Ahel, I. Fragment Binding to the Nsp3 Macrodomain of SARS-CoV-2 Identified through Crystallographic Screening and Computational Docking. *Science Advances* **2021**, *7*, eabf8711.

(S68) Wojdyr, M. GEMMI: A Library for Structural Biology. *Journal of Open Source Software* **2022**, *7*, 4200.

(S69) deSolms, S. J.; Ciccarone, T.; MacTough, S.; Shaw, A. W.; Buser, C.; Ellis-Hutchings, M.; Fernandes, C.; Hamilton, K.; Huber, H.; Kohl, N.; Lobell, R.; Robinson, R.; Tsou, N.; Walsh, E.; Graham, S.; Beese, L.; Taylor, J. Co-Crystal Structure

Of Human Farnesyltransferase With Farnesyldiphosphate and Inhibitor Compound 33a. 2002; https://doi.org/10.2210/pdb1MZC/pdb.

(S70) deSolms, S. J.; Ciccarone, T. M.; MacTough, S. C.; Shaw, A. W.; Buser, C. A.; Ellis-Hutchings, M.; Fernandes, C.; Hamilton, K. A.; Huber, H. E.; Kohl, N. E.; Lobell, R. B.; Robinson, R. G.; Tsou, N. N.; Walsh, E. S.; Graham, S. L.; Beese, L. S.; Taylor, J. S. Dual Protein Farnesyltransferase-Geranylgeranyltransferase-I Inhibitors as Potential Cancer Chemotherapeutic Agents. *Journal of Medicinal Chemistry* **2003**, *46*, 2973–2984.

(S71) Pavlicek, J.; Ptacek, J.; Cerny, J.; Byun, Y.; Skultetyova, L.; Pomper, M.; Lubkowski, J.; Barinka, C. X-Ray Structure of of Human Glutamate Carboxypeptidase II (GCPII) in a Complex with CCIBzL, a Urea-Based Inhibitor N~2~-[(1-Carboxycyclopropyl)Carbamoyl]-N~6~-(4-Iodobenzoyl)-L-lysine: 4oc0. 2014; https://www.wwpdb.org/pdb?id=pdb_00004oc0.

(S72) Pavlicek, J.; Ptacek, J.; Cerny, J.; Byun, Y.; Skultetyova, L.; Pomper, M. G.; Lubkowski, J.; Barinka, C. Structural Characterization of P1′-Diversified Urea-Based Inhibitors of Glutamate Carboxypeptidase II. *Bioorganic & Medicinal Chemistry Letters* **2014**, *24*, 2340–2345.

(S73) Fährrolfes, R.; Bietz, S.; Flachsenberg, F.; Meyder, A.; Nittinger, E.; Otto, T.; Volkamer, A.; Rarey, M. Proteins Plus: A Web Portal for Structure Analysis of Macromolecules. *Nucleic Acids Research* **2017**, *45*, W337–W343.

(S74) Schöning-Stierand, K.; Diedrich, K.; Fährrolfes, R.; Flachsenberg, F.; Meyder, A.; Nittinger, E.; Steinegger, R.; Rarey, M. Proteins Plus: Interactive Analysis of Protein–Ligand Binding Interfaces. *Nucleic Acids Research* **2020**, *48*, W48–W53.

(S75) Schöning-Stierand, K.; Diedrich, K.; Ehrt, C.; Flachsenberg, F.; Graef, J.; Sieg, J.; Penner, P.; Poppinga, M.; Ungethüm, A.; Rarey, M. Proteins Plus: A Comprehensive

Collection of Web-Based Molecular Modeling Tools. *Nucleic Acids Research* **2022**, *50*, W611–W615.

(S76) Mysinger, M. M.; Carchia, M.; Irwin, J. J.; Shoichet, B. K. Directory of Useful Decoys, Enhanced (DUD-E): Better Ligands and Decoys for Better Benchmarking. *Journal of Medicinal Chemistry* **2012**, *55*, 6582–6594.

(S77) Suino-Powell, K.; Xu, Y.; Zhang, C.; Tao, Y.-g.; Tolbert, W. D.; Simons, S. S.; Xu, H. E. Doubling the Size of the Glucocorticoid Receptor Ligand Binding Pocket by Deacylcortivazol. *Molecular and Cellular Biology* **2008**, *28*, 1915–1923.

(S78) Xu, H. Doubling the Size of the Glucocorticoid Receptor Ligand Binding Pocket by Deacylcortivazol: 3bqd. 2008; `https://www.wwpdb.org/pdb?id=pdb_00003bqd`.

(S79) Sobolevsky, A.; Rosconi, M.; Gouaux, E. Isolated Ligand Binding Domain Dimer of GluA2 Ionotropic Glutamate Receptor in Complex with Glutamate, LY 404187 and ZK 200775: 3kgc. 2009; `https://www.wwpdb.org/pdb?id=pdb_00003kgc`.

(S80) Hast, M. A.; Fletcher, S.; Cummings, C. G.; Pusateri, E. E.; Blaskovich, M. A.; Rivas, K.; Gelb, M. H.; Van Voorhis, W. C.; Sebti, S. M.; Hamilton, A. D.; Beese, L. S. Structural Basis for Binding and Selectivity of Antimalarial and Anticancer Ethylenediamine Inhibitors to Protein Farnesyltransferase. *Chemistry & Biology* **2009**, *16*, 181–192.

(S81) Hast, M.; Beese, L. Protein Farnesyltransferase Complexed with Bisubstrate Ethylenediamine Scaffold Inhibitor 5. 2009; `https://doi.org/10.2210/pdb1E37/pdb`.

(S82) Lippa, B.; Pan, G.; Corbett, M.; Li, C.; Kauffman, G. S.; Pandit, J.; Robinson, S.; Wei, L.; Kozina, E.; Marr, E. S.; Borzillo, G.; Knauth, E.; Barbacci-Tobin, E. G.; Vincent, P.; Troutman, M.; Baker, D.; Rajamohan, F.; Kakar, S.; Clark, T.; Morris, J. Synthesis and Structure Based Optimization of Novel Akt Inhibitors. *Bioorganic & Medicinal Chemistry Letters* **2008**, *18*, 3359–3363.

(S83) Pandit, J. Crystal Structure of Akt-1 Complexed with Substrate Peptide and Inhibitor: 3cqw. 2008; https://www.wwpdb.org/pdb?id=pdb_00003cqw.

(S84) Boettcher, J.; Specker, E.; Heine, A.; Klebe, G. HIV-1 Protease in Complex with Pyrrolidinmethanamine: 1xl2. 2005; https://www.wwpdb.org/pdb?id=pdb_00001xl2.

(S85) Specker, E.; Böttcher, J.; Lilie, H.; Heine, A.; Schoop, A.; Müller, G.; Griebenow, N.; Klebe, G. An Old Target Revisited: Two New Privileged Skeletons and an Unexpected Binding Mode For HIV-Protease Inhibitors. *Angewandte Chemie International Edition* **2005**, *44*, 3140–3144.

(S86) Peat, T.; Newman, J.; Deadman, J.; Rhodes, D. Structural Basis for a New Mechanism of Inhibition of HIV Integrase Identified by Fragment Screening and Structure Based Design: 3nf7. 2011; https://www.wwpdb.org/pdb?id=pdb_00003nf7.

(S87) Rhodes, D. I.; Peat, T. S.; Vandegraaff, N.; Jeevarajah, D.; Le, G.; Jones, E. D.; Smith, J. A.; Coates, J. A.; Winfeld, L. J.; Thienthong, N.; Newman, J.; Lucent, D.; Ryan, J. H.; Savage, G. P.; Francis, C. L.; Deadman, J. J. Structural Basis for a New Mechanism of Inhibition of H I V-1 Integrase Identified by Fragment Screening and Structure-Based Design. *Antiviral Chemistry and Chemotherapy* **2011**, *21*, 155–168.
